# Nausea reshapes visual exploration and pupillary responses to affective images

**DOI:** 10.64898/2026.07.17.739163

**Authors:** Irene Petrizzo, Fadila Hadj-Bouziane, Elvio Blini

## Abstract

Perception is a continuous, active, and dynamic process of inference shaped by current goals, past experience, and hard biological constraints. The physiological state of the body is arguably the most fundamental one. Continuous inference about interoceptive states colors perception with emotions, which in turn affect basic perceptual and attentional processes. How, then, do we *really* look at the world when experiencing malaise?

Here we leverage Galvanic Vestibular Stimulation (GVS) – a non-invasive brain stimulation technique that induces mild dizziness, motion sickness, and nausea – in conjunction with high-resolution eye tracking, pupillometry, and computational modelling. We demonstrate that mild vestibular perturbations disrupt the coupling between gaze and informative low-level visual features. Crucially, however, this effect depends on individual susceptibility and on stimulus valence. Particularly susceptible participants experiencing stronger nausea *increase* their reliance on visually salient features, and particularly so for threatening (negatively valenced) images. Pupillometry reveals larger tonic pupil size in these participants, consistent with heightened arousal, as well as disrupted phasic responses to affective pictures.

Altogether, these results unveil the continuous and dynamic interplay between interoceptive and exteroceptive processes, showing that how we perceive the environment fundamentally resonates with how we feel.

**Significance statement:** Feeling unwell – whether from illness or motion – is a universal experience. Bodily signals profoundly shape cognition, emotion, and decision-making. We therefore asked how nausea changes visual exploration of the environment. We electrically stimulated the vestibular system of healthy volunteers to induce dizziness and nausea, while monitoring gaze during free viewing of affective images. The stronger the nausea, the more visual exploration was guided by objectively salient visual features, especially for highly arousing images. Pupillometry revealed larger pupils, consistent with heightened arousal, and disrupted pupillary responses to affective pictures, showing less room for affective processing and increased cognitive control. We conclude that nausea may dull perception, while strengthening its grip on physical reality.

**Highlights:**

- GVS caused variable degrees of dizziness and nausea
- GVS led to spatial biases in visual exploration which were, however, further dependent on image valence and content
- During GVS eye fixations landed, on average, at locations that were less salient based on their mere physical properties, provided participants did not experience strong interoceptive perturbations
- Highly susceptible participants showed the opposite pattern: their fixations landed on visually more salient and informative locations, especially for negative images
- Participants who felt stronger effects, including nausea, also showed the largest increase in tonic pupil size; concurrently, the signature phasic dilation associated with negative emotions was disrupted
- Oculomotor behavior and perception adjust dynamically to ongoing interoceptive processes; GVS can be an effective tool to study this dialogue, provided that inter-individual differences are accounted for

## Introduction

> *“The Nausea is not inside me: I feel it out there in the wall, in the suspenders, everywhere around me.*
>
> *It makes itself one with the café, I am the one who is within it.”*
>
> *(Jean-Paul Sartre)*

The boundaries between interoception, sensing our body, and exteroception, sensing the world, are blurry (Toussaint et al., 2024); the case of vestibular perception is a particularly compelling example. Traditionally, interoception encompasses the afferent processing of visceral signals originating within the body and which refer to its current state (Craig, 2002; Critchley & Garfinkel, 2017). The vestibular system is instead designed to detect gravitational and inertial forces (linear and angular acceleration of the head) through an ensemble of five sensory organs located in the inner ear, which places it in the realm of proprioception or exteroception. However, recent framings of interoception are more inclusive (Blini et al., 2020; Ceunen et al., 2016; Quadt et al., 2018; Toussaint et al., 2024; Tsakiris & Critchley, 2016), using it as an umbrella term for the phenomenological experience of the body state, regardless of which input informs that experience.

Multisensory interoceptive signals are increasingly recognized to play an important, pervasive role in several aspects of cognition, emotion, and decision-making, in health and disease (Critchley & Garfinkel, 2017; Quadt et al., 2018; Tsakiris & Critchley, 2016). A similar shift from low-level (e.g., oculo-motor adjustments) to higher-level processing, including affect (Carmona et al., 2009; Deroualle & Lopez, 2014; Lopez, 2016; Mast et al., 2014), has been documented for the vestibular sense (Deroualle & Lopez, 2014; Ferrè & Haggard, 2020). The vestibular system is represented, at the cortical level, in a highly-distributed network of inherently multisensory areas, chiefly centered around the so-called parieto-insular vestibular cortex (Angelaki et al., 2009; Lopez et al., 2012; zu Eulenburg et al., 2012). This (multisensory) feature firmly places the vestibular system in a privileged position to provide a universal grounding for bodily self-consciousness (Lenggenhager & Lopez, 2015; Pfeiffer et al., 2014; Tsakiris, 2017). Moreover, it provides a working model to study the dialogue between multisensory interoception and basic perceptual and affective processes. This point is made clearer when considering how the universal human experience of feeling unwell – whether from illness, fatigue, or motion – can fundamentally reshape homeostatic priorities, and thus motivated behavior and mood (Blini et al., 2018; Preuss et al., 2014). This interaction is framed more formally by predictive processing accounts (Barrett & Simmons, 2015; Seth, 2013). The brain would continuously generate predictions about the causes of both interoceptive and exteroceptive signals, using the actual sensory input primarily to finely-tune these inferences (Barrett & Simmons, 2015). Conscious perception emerges from the integration of predictions and prediction errors across interoceptive and exteroceptive domains, with interoception playing a central role in shaping affective experience (Seth, 2013). From this perspective, interoceptive perturbations are not merely the passive consequence of altered sensory signals, but a bias of the brain’s inferential regime which can, in turn, propagate to how external information is sampled, weighted, and ultimately perceived (Blini et al., 2020).

Our understanding of interoceptive processes is limited by the difficulties in causally manipulating interoceptive states (Tsakiris, 2017). Using “interoception” in its broader sense, however, enables much more methodological flexibility. The vestibular system, for instance, can be experimentally manipulated precisely by thermal or electrical means. Galvanic Vestibular Stimulation (GVS) involves weak electrical currents applied to the skin over the mastoid processes (behind the ears): GVS targets the VIII cranial nerve and thus modulates both otolith and semicircular canal afferents (Fitzpatrick & Day, 2004; Kwan et al., 2019). At the behavioral level, the result is a torsional movement of the eyes (Marchand et al., 2025) alongside illusions of body movement, dizziness, and a variable degree of motion sickness and nausea (Hammam et al., 2012; Quinn et al., 2015; Yates et al., 2014). It is well established that GVS can bias spatial attention and perception (Angelaki et al., 2009; Ferrè et al., 2013; Lenggenhager et al., 2007), largely due to hemispheric asymmetries in the targeted brain regions. Vestibular stimulation is indeed a very effective tool to ameliorate lateralized disorders of spatial awareness, such as spatial neglect (Cappa et al., 1987; Karnath et al., 1996; Rode et al., 1995), and related manifestations (e.g., somatoparaphrenia, a body-related delusion Bisiach et al., 1991; Rode et al., 1992).

The effects of vestibular stimulation, however, also extend beyond spatial attention, influencing body representation, ownership, and awareness (Ferrè & Haggard, 2016; Hoover & Harris, 2015; Lopez et al., 2010; Ponzo et al., 2018). For example, the rubber-hand illusion (Botvinick & Cohen, 1998) is an established experimental model of bodily awareness. In the typical setting, the participant’s hand is hidden from view while a fake hand is placed in sight (in a plausible position); then, both are stroked synchronously with a brush. The temporal and spatial congruence leads the participants to experience the fake hand as their own. Essentially, the rubber-hand paradigm probes the permeable interface between exteroceptive signals and body-related representations. This is often measured through the shift in perceived hand location (proprioceptive drift), a measure of visual capture strength by the fake hand (Pavani et al., 2000). Conflicting results have emerged when GVS has been applied in the context of the rubber hand illusion. Ferrè et al. (2015), for example, reported decreased visual capture by the rubber hand with GVS, whereas other authors reported an increase (Lopez et al., 2010; Ponzo et al., 2018). The incongruence has been chiefly ascribed to methodological differences, most notably the duration of the electric stimulation. The modulation of visual capture (regardless of its direction) has been interpreted, in keeping with predictive coding accounts, in terms of GVS inducing a profound reweighting of low-level sensory signals in the process of multisensory integration, with the weights assigned to vision suggested to either decrease (Ferrè et al., 2015) or increase (Ponzo et al., 2018). The emergence of the rubber-hand illusion itself is attributed to decreased precision of sensory input (e.g., somatosensory), prompting top-down inferences to resolve the conflict (Zeller et al., 2015). Interestingly, affective valence (i.e., affective touch) may further modulate this reweighting process during GVS (Ponzo et al., 2018), indicating a complex interaction between interoceptive signals, affect, and sensory processing.

Here we take a more radical reductionist approach. Instead of focusing on multisensory settings, we isolate the visual modality. However, we move beyond the more traditional metrics of visual exploration – e.g., dwell time, fixation location – to assess more elusive properties of gaze such as the reliance on bottom-up (physical) features vs top-down (endogenous) processes. In order to do so, we also move beyond classic laboratory settings, that involve simple to absent visual stimulation when recording eye movements, to present instead complex, and relatively more naturalistic image stimuli. While GVS is known to induce systematic shifts of attention and eye movements, its impact on the spontaneous visual exploration of the environment is relatively unexplored. We reasoned that GVS and interoceptive perturbations may reshape, prior to multisensory processes, also basic unisensory (visual) processes beyond these “reflexive” shifts, but in an *adaptive* and context-dependent manner (e.g., differing by valence).

Several computational approaches have characterized image features according to objective properties (e.g., luminance, color, intensity; Harel et al., 2007; Itti & Koch, 2000). The predictions of these saliency maps correlate well with spontaneous eye movements during free viewing. However, an “affective gap” has been described consisting of decreased adherence to stimulus-driven salience when the overall emotional valence prevails (Niu et al., 2012; Pilarczyk et al., 2021). This approach therefore probes the dynamic balance between bottom-up and top-down processes and can be leveraged to indirectly assess reliance on the visual modality; the assumption is that more physically salient features also offer more stable and informative cues to the visual system, which is arguably a priority under sensory conflict and vestibular perturbations specifically.

We therefore administered GVS (vs sham) stimulation to 42 healthy participants, in separate sessions (**Figure 1**). We choose stimulation parameters that are known to elicit reliable motion sickness, though at the group level. Within each session, furthermore, we intermixed stimulation and no stimulation trials (online vs offline), which provided within-session control conditions. Participants were presented with images chosen from an established normative database (Kurdi et al., 2017) and selected for their valence (negative, neutral, positive images). High-resolution eye tracking was in place to monitor the participants’ gaze continuously during free viewing. There were, however, two separate tasks, optimized for different questions. In the image viewing task, visual exploration was fully unconstrained; it was used to characterize spontaneous oculomotor behavior. In the second task, we constrained gaze to a narrow central spot to better assess changes in pupil size.

**Figure 1.**
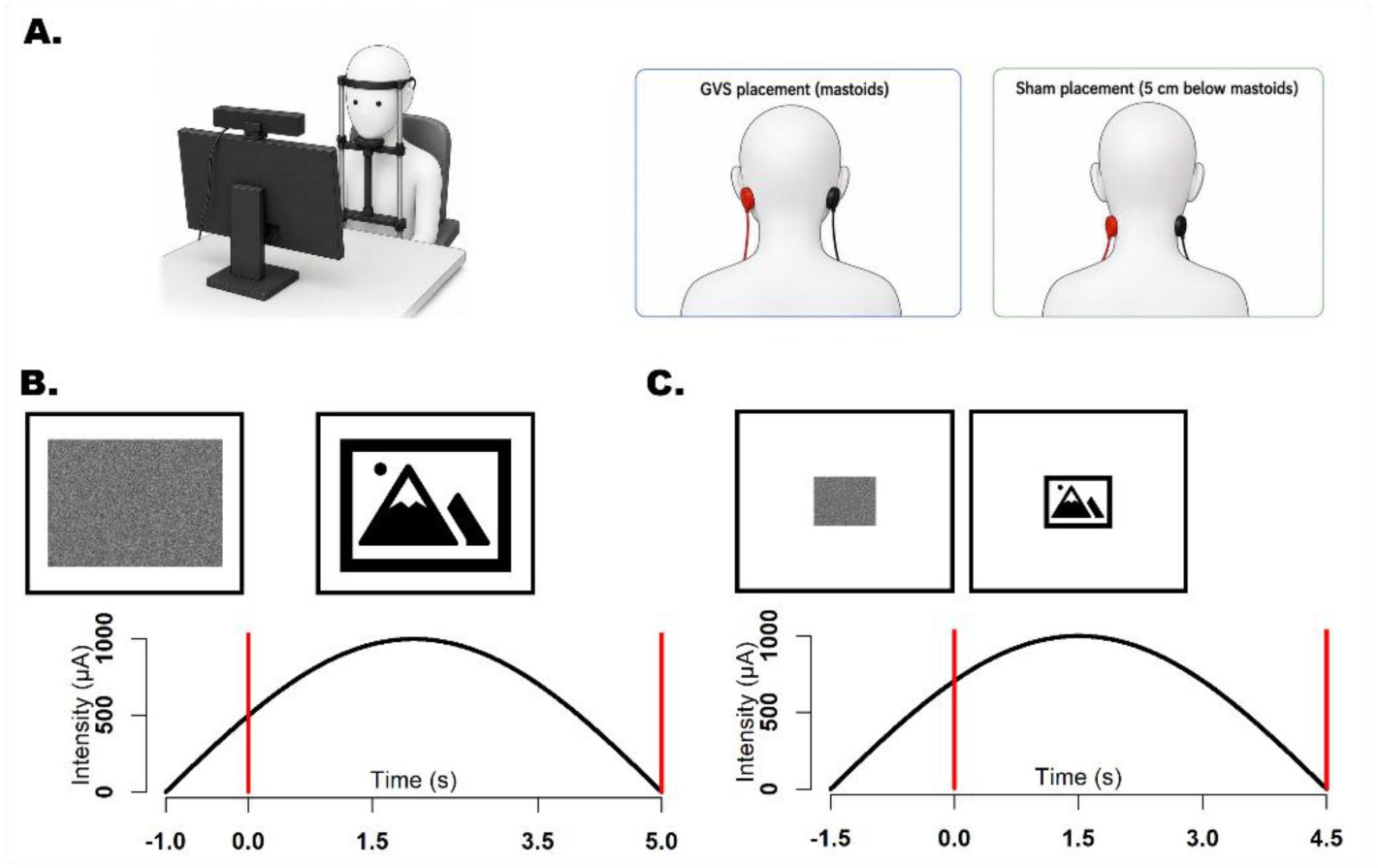
Methods and procedures. **A. Vestibular stimulation.** Electric stimulation (left anode, right cathode) was delivered either on the mastoids, to stimulate the vestibular system (L-GVS), or on the neck (sham), in different sessions. Electric pulses were delivered as low-frequency half-sinusoidal waves synchronized with the presentation of visual stimuli. Each pulse had a total duration of 6s and reached a peak intensity of 1mA. In both sessions, the stimulation was delivered on only half of the trials to probe online vs offline effects. A high-resolution eye tracker monitored the participants’ gaze continuously. Visual stimuli were images drawn from a standardized database, divided into three different valence categories: negative, neutral, and positive affect images. Panel A has been generated with the assistance of a LLM (GPT-5.5). **B. Image Viewing Task.** Each target image (36x26°) was presented on screen for 5s. Participants were asked to freely explore the image. The main measures of interest were the distribution of the fixations (center of gravity), and the physical saliency associated with gaze coordinates. **C. Pupillometry task.** Visual stimuli were in this case 7x5.5° grayscale images with the same average luminance. Participants were asked to maintain central fixation throughout the task to accurately measure pupil size, in both its tonic and phasic components.

The measurement of pupil size gained significant traction and renewed interest in recent years as a non-invasive tool to probe autonomic balance (Binda & Gamlin, 2017; Castellotti, Castaldi, et al., 2025); as such it is perfectly suited to probe affective and emotional responses to visual stimuli (Bogdanova et al., 2022; Dureux et al., 2021; Gilzenrat et al., 2010; Murphy et al., 2011). Altogether, these manipulations were designed to respond to the following question: how do we look at the world when experiencing nausea, and how do we see the emotions therein?

## Materials and methods

### Participants and recruiting plan

The rationale and computer code for the power analysis and recruiting plan are outlined at length in **Supplementary 1**. The main question was whether GVS impacts visual exploration and, if so, whether the effect is context-sensitive (i.e., it depends on image valence). A few studies explored the impact of GVS on the weighting of visual information during multisensory tasks, notably the rubber-hand illusion (Ferrè et al., 2015; Lopez et al., 2010; Ponzo et al., 2018). This is a very different, high-level paradigm compared to our choice of using a more direct, purely visual task (i.e., image viewing and eye tracking). Furthermore, these studies present discordant results, chiefly ascribed to key differences in the stimulation parameters (Ferrè et al., 2015; Lopez, 2015, 2016) which we hereby perpetuate (see below). Interestingly, one study also explored the effect of valence by delivering the tactile stimulations that determine the rubber-hand illusion at different speeds (CT-optimal, i.e. affective touch, or not) and found a three-way interaction between session (sham, GVS), synchrony (synchronous, asynchronous stimulation), and speed (Ponzo et al., 2018). This setting is conceptually similar to our design, which consists of session (sham, GVS), stimulation (on, off), and valence (in our case positive, neutral, and negative images, see below). Ponzo et al. (2018) report an effect size (partial eta squared) of roughly 0.193. A power analysis based on a general F test suggests that 90% statistical power for this effect size and design (2x2x3, fully within) would be achieved after testing 28 participants (alpha= 0.05). We decided to enroll a maximum of 42 participants, which ensures good statistical power (>80%) for substantially smaller effect sizes (**Supplementary 1**). However, we also planned interim analyses at N= 30. As a result, the significance threshold alpha that preserves a cumulative 5% error rate across the experiment is 0.034 (Pocock, 1977).

Forty-five participants met the inclusion and exclusion criteria. The inclusion criteria were right-handedness (as assessed by the Edinburgh handedness inventory (Oldfield, 1971) and normal or corrected-to-normal vision. The exclusion criteria included neurologic, psychiatric, cardiac, or otologic disorders, as well as the presence of metallic implants or splinters in the body or skin conditions in the neck and head regions. Three participants were excluded from the study: calibration of the eye tracker was impossible in one case; two participants withdrew their participation from the study for excessive eye fatigue and for feeling unwell. The final sample was composed of 42 young adults (mean age: 23.5 years, SD: 6 years), in majority women (N= 28, 66%). The participants were mostly students at the University of Trento and attended for either course credits or a small monetary fee; they all provided their written informed consent for the inclusion. The study was approved by the ethics committee of the University of Trento (Comitato Etico per la Ricerca) with protocol number 2025-046 and adhered to the guidelines laid out by the Declaration of Helsinki.

### Procedures

#### Galvanic Vestibular Stimulation (GVS)

GVS was delivered via a commercial brain stimulator (BrainStim, EMS, Bologna). We used a bilateral montage with the anode on the left mastoid process, and the cathode on the right mastoid process (known as L-GVS). L-GVS mainly activates right hemisphere structures (Lopez et al., 2012; zu Eulenburg et al., 2012), and has been found particularly effective in modulating the degree of visual capture in multisensory tasks (Ferrè et al., 2015; Lopez et al., 2010; Ponzo et al., 2018). A sham condition was also included, with electrodes placed symmetrically (the anode on the left) about 5 cm below the mastoids, above the neck, and distant from the trapezoidal muscles yielding proprioceptive signals (Lenggenhager et al., 2007). The two different electrode positions are depicted in Figure 1A. GVS and sham were delivered in two separate sessions. In both sessions, the area of application was first cleaned with abrasive gel (Abralyt HiCl). Two circular rubber electrodes (3 cm diameter) were then fixed in position with conductive paste (Ten20) and adhesive tape.

The application of small current intensities over the mastoid bones is, generally speaking, well tolerated (Utz et al., 2011). Vestibular specific effects include, however, the illusion of head and body movements, resulting in transient dizziness. Nausea is per se rare or mild, tolerable, but stimulation parameters are known to play a critical role in this regard. While continuous stimulation results in rapid habituation, sinusoidal waveforms optimally activate both otolith and semicircular canal afferents (Kwan et al., 2019), particularly at high frequencies. On the other hand, low frequencies modulate more effectively skin sympathetic nerve activity and evoke motion sickness and nausea more frequently, likely due to sensory mismatches (Hammam et al., 2012; Quinn et al., 2015; Yates et al., 2014). Here we decided to use low-frequency half-sinusoidal waveforms so that we could effectively exploit this feature and at the same time induce the tonic polarity unbalance that leads to increased right hemisphere activation. The electric pulses lasted 6 seconds each (hence ≈0.08 Hz if considering the full cycle); the current intensity reached a fixed 1mA peak well within the active part of the behavioral tasks described below (see **Figure 1**). We decided to use the same parameters for both GVS and sham conditions to control very tightly the unspecific effects of electrical stimulation (e.g., itching, tingling, burning, discomfort), which are very likely to result in an increased state of arousal. The obvious downside is that participants are potentially able to tell sham from GVS conditions based on the position of the electrodes – though this aspect was evaluated informally at debriefing and did not emerge as evident in participants’ reports. Within both GVS and sham, furthermore, the stimulation was randomly intermixed with no stimulation trials (50% probability). This additional control was intended to assess the immediate effect of expectations – that is, the anticipation of an electrical stimulation, which also constitutes an important arousing factor (Finke et al., 2021). It should be stressed, however, that the fast pacing of the tasks implies that no stimulation trials are likely affected, particularly during GVS, by long-term changes in brain activation driven by plasticity as well as by specific behavioral aftereffects (such as persistent motion sickness following the end of electrical pulses).

The stimulation was delivered only after an initial impedance check. Prior to each session, participants were given a few trains of stimulation to familiarize themselves with the resulting sensations. The order of GVS and sham conditions was counterbalanced across participants. In between the two sessions, participants took a long break and were invited to walk freely, thereby minimizing potential carryover effects. At the end of each stimulation session, a brief questionnaire was administered to participants to gather their overall subjective sensations. We evaluated the following aspects: 1) intensity of the itching/tingling sensation caused by the electrical currents; 2) the degree of movement illusions/dizziness; 3) the amount of nausea/motion sickness; 4) the overall pain/discomfort throughout the session. Participants rated their sensations on a scale from 1 (not at all) to 10 (extreme).

#### Eye-tracking

Participants were tested in a dimly lit, quiet room. They were instructed to rest their head comfortably on the chinrest of an infrared-based eye tracker (EyeLink 1000 Tower Mount, SR research Ltd.). Each task started with the built-in calibration procedure, after which the eye tracker monitored gaze and pupil size continuously at 1000 Hz. The open-source software OpenSesame (Mathôt et al., 2012) was used to display experimental stimuli on the screen and collect the data. All participants performed two Sessions (GVS or sham, counterbalanced). In each session they performed two tasks in a fixed order: the Image Viewing task (IVT) and the Pupillometry task (PT) both consisted of passive viewing of images but were optimized respectively for the analysis of free oculomotor exploration and pupil size.

#### Image-Viewing task (IVT)

The image viewing task closely aligns with the timings and parameters of a previous study (Cinetto et al., 2025), except that stimuli were selected by their valence (see paragraph below). Each trial started with the presentation of a noisy image, which acted as a break in between different images. In half of the trials, in both GVS and sham sessions, the electrical stimulation started during this noisy-image phase (1 s) and then continued, peaked, and ended during image presentation. The subsequent target images lasted on screen 5 s (Cinetto et al., 2025), see **Figure 1B**. All images in this task covered most of the screen, which was located at roughly 46 cm distance from participants, and encompassed about 32 x 26 degrees of visual angle. There were 36 different images, selected from three different categories (Negative, Neutral, Positive valence). Each image was presented twice (Stimulation ON vs OFF), in a fully randomized order, totaling 72 trials per Session (GVS vs sham) and 144 total. The task was divided into 2 blocks of equal length, which provided the opportunity for participants to rest their eyes.

We have decided to exploit the increased statistical power of a within-subjects design in this work, which implies repeated image exposure. Repeated image exposure can alter eye-movement patterns, but nevertheless individuals tend to revisit similar regions and remain attracted by salient features (Kaspar & König, 2011a, 2011b). Therefore, repeated image exposure appears to be a viable choice if the aim is to assess whether GVS disrupts this otherwise reliable tendency. We have mitigated potential drawbacks by counterbalancing GVS and sham conditions.

#### Pupillometry task (PT)

The pupillometry task is similar to that presented above, though several parameters have been optimized for the measurement of pupil size. First, the images and noise mask were much smaller (7 x 5.5 °) to constrain eye movements, which may bias the recording of pupil size. Second, unlike the IVT, all stimuli were transformed into grayscale images, and were further processed in order to have the same mean luminance following the procedure of . Third, we increased the inter trial interval and timings to cope with the fact that pupil size is a rather slow physiological signal (Blini et al., 2024; Castellotti, Petrizzo, et al., 2025) and requires time to reach stable baseline levels following visual stimulations. In particular, the noisy image lasted on screen a minimum of 2 s (randomly jittered up to 3 s). During ON trials, the electric stimulation started 1.5s before stimulus presentation and then peaked and ended during image presentation (4.5 s), see **Figure 1C**. Target images were made of the same 36 exemplars used for the IVT. They belonged to three different categories (Negative, Neutral, Positive valence). Each image was presented with or without electrical stimulation (50% probability) a total of 6 times, totaling 216 trials per Session (GVS vs sham) and 432 overall. Image presentation was fully randomized. Due to the length of this task, the PT was divided into 3 blocks of equal length, which provided the opportunity for participants to rest their eyes.

#### Stimuli selection and subjective ratings

The target images were selected from the Open Affective Standardized Image Set (OASIS) (Kurdi et al., 2017). We limited our selection to images depicting people, because we assumed them behaviorally more salient and internally more homogeneous. The database includes a relatively recent (2015) standardization and normative data (N= 822) for two fundamental dimensions: valence – the degree of positive or negative affective response – and arousal – the intensity of the affective response. The final selection consisted of 36 images, forming three clearly distinct categories (12 images each). Negative images had on average low valence scores (M= 1.85, SD= 0.67) and high arousal scores (M= 4.39, SD= 0.56). Positive images had on average both high valence scores (M= 5.69, SD= 0.29) and high arousal scores (M= 4.41, SD= 0.43). Neutral images had on average medium valence scores (M= 4.44, SD= 0.51) and low arousal scores (M= 3.17, SD= 0.66). The normative data and statistics of the stimuli are reported in **Supplementary 2**. In short, the three categories were all clearly separable in terms of valence; negative and positive images, furthermore, presented higher arousal scores than the neutral category, but did not differ from each other. This is important because some of the measures of this study, i.e. pupil size, are known to respond more prominently to the arousal dimension, which is why we strived to match the two affective categories in this regard.

However, we were also interested in the participants’ subjective evaluation of the images. For this reason, all participants, after the main experiments, individually rated each image on the two dimensions. The rating procedure differs from the original for a few key details. Valence and arousal were rated independently in the normative data to avoid contamination; we were, however, interested in both dimensions, and thus cannot rule out potential carryover effects, since we asked participants to rate arousal and valence sequentially. Furthermore, we adapted the question for the arousal dimension so that it focused on the physiological impact of the images (from 1 – very calming – to 10 – very activating). We thought this framing was more directly relevant for the analysis of physiological signals such as pupil size.

### Data Preprocessing

The data were analyzed with R 4.2.3 (R Core Team, 2023). The following packages substantially improved our workflow: *buildmer* (Voeten, 2019); *lme4* ; *Pupilla* (Castellotti, Petrizzo, et al., 2025); *emmeans* (Lenth, 2019) ; *saccades* (Malsburg, 2019); *afex* (Singmann et al., 2023); *ggplot2* (Wickham, 2016); *dplyr* (Wickham et al., 2023). All materials, raw data, and computer code are available without restrictions at the following link: https://osf.io/dwjzr

#### Image-Viewing task (IVT)

The continuous stream of gaze samples was first segmented into stable fixation events. This was accomplished via a well-established, velocity-based criterion (Malsburg, 2019). The algorithm considers the velocity distribution for each trial and classifies as saccades the displacements exceeding +6 standard deviations of that distribution (Cinetto et al., 2025; Engbert & Kliegl, 2003). We then discarded fixations falling outside the borders of the image and, to avoid the inclusion of spurious events, fixations shorter than 100 ms. Several parameters were then computed at the fixation level: the center of gravity (both on the horizontal and vertical axes); the within-fixation dispersion of the samples, i.e. their median absolute deviation (both on the horizontal and vertical axes, then pooled to obtain a single value); their temporal duration. The fixations associated with a large uncertainty (dispersion > 2 standard deviations) were discarded.

It was particularly important, in the context of this work, to move beyond basic oculomotor parameters to explore the reliance on specific elements of each image. Different elements depicted in a scene can vary for their intrinsic interest as well as for their physical properties. The latter feature can be quantified precisely – albeit in a model-driven fashion – and can be exploited to create a visual saliency map for a target image. Here, we used the Graph-Based Visual Saliency model (Harel et al., 2007) to compute a saliency map for each of the 36 images. The maps are obtained by first calculating activation maps for several physical features (e.g., color, orientation, intensity); the features are then linearly combined in a final saliency map with normalized values (0, least salient, to 1, most salient) (Reynaud et al., 2021). Therefore, for each fixation we retrieved the corresponding saliency score. The saliency score was taken in correspondence of the fixation’s center of gravity. However, to avoid overfitting we also considered a window of a few pixels around the center of gravity and took the average saliency score of all pixels in the window. The natural choice was using the median absolute deviation of the coordinates of all samples composing a fixation as a measure of dispersion and thus size of the window (the larger the dispersion, the larger the window). However, in control analyses we kept the size of the window fixed to either low values (6x6 pixels) or high values (10x10 pixels). The saliency scores calculated this way were highly correlated (r> 0.99) likely due to the smoothing applied to the saliency maps. We therefore present the data for visual saliency computed in a window depending on each fixation’s dispersion.

There are several approaches to compare ground-truth fixation patterns and saliency maps (Le Meur & Baccino, 2013). In addition to comparing the saliency associated with each fixation, one particularly robust method requires comparing saliency maps and fixation-density maps directly via a given similarity measure (e.g., Pearson’s correlation). This approach has the only drawback that reliable fixation-density maps can only be obtained by collapsing all participants together, since a 5 s time window for the exploration of images generally includes too few fixations. However, we also followed this strategy to ensure consistency in the main results.

Finally, fixation-wise scores were complemented by trial-wise scores, namely the overall number of fixations, the average fixation duration, and dwell time, computed as the overall time spent fixating the image (thus excluding blinks, saccades, or artifacts). All these variables were submitted to statistical modelling.

Overall, data preprocessing for this task led to minimal data loss (on average 3.2 % of trials discarded per participant, SD= 5.9%).

#### Pupillometry task (PT)

Pupil size was recorded from the left eye, in arbitrary units of area (a.u.). We retained only the values measured during fixation events, as identified by the eye tracker (i.e., measures during blinks and saccades were discarded). We then removed values below 2.5 standard deviations from each participant’s average pupil size and used a velocity-based criterion to remove implausibly fast changes in pupil size, as well as 120 ms of recordings around these values. These procedures were adopted to minimize the impact of artifacts such as partial occlusion of the pupil by the eyelid. Trials in which more than 40% of the data were missing were discarded. The gaps in the remaining trials were linearly interpolated, then traces were smoothed with cubic splines. Finally, traces were down-sampled to 20-ms epochs by taking the median pupil size within each time bin. The preprocessing pipeline largely follows previous studies (Blini et al., 2024) and the default parameters of the R library *Pupilla* (Castellotti, Petrizzo, et al., 2025). Overall, data cleaning procedures led to discarding on average 7.5% of trials per participant (SD= 10%).

We deviate from previous studies in that here we do not present z-transformed pupil size data. We found that the type of stimulation has a remarkable effect on tonic pupil size: GVS was associated with significantly larger baseline pupil size. We therefore present the data in arbitrary eye tracker units (a.u.) to avoid potential distortions, although the results hold when z-scoring the data both i) separately and ii) across conditions. Finally, all series were realigned by subtraction to trial-wise baseline values which varied as a function of the experimental question. We used the time window from -1600 ms to -1500 ms as baseline to study the effect of electrical stimulation (both GVS or sham). We used the time window from -100 ms to 0 ms instead to better assess the perceptual and affective processing of images.

## Results

### GVS induces a variable degree of dizziness and motion sickness

After each session, we asked participants to rate their experience across four dimensions on a scale from 1 to 10. We took their subjective reports for the sham stimulation and subtracted them from GVS reports, thus having positive values when a given dimension was felt more clearly during GVS. As depicted in **Figure 2A**, this was the case for illusions of movement and dizziness, which were reported by most participants (32/42, 76.2%) and were significant at the group level (*t*(41)= 7.22, *p*< 0.001). Nausea and motion sickness were also reported by a smaller subset of participants (18/42, 42.9%) – all but one of whom also reported illusions of movement – and were significant at the group level (*t*(41)= 4.01, *p*< 0.001). There was a moderate positive correlation between illusions of movement and motion sickness (r= 0.44, *t*(40)= 3.14, *p*= 0.003).

**Figure 2.**
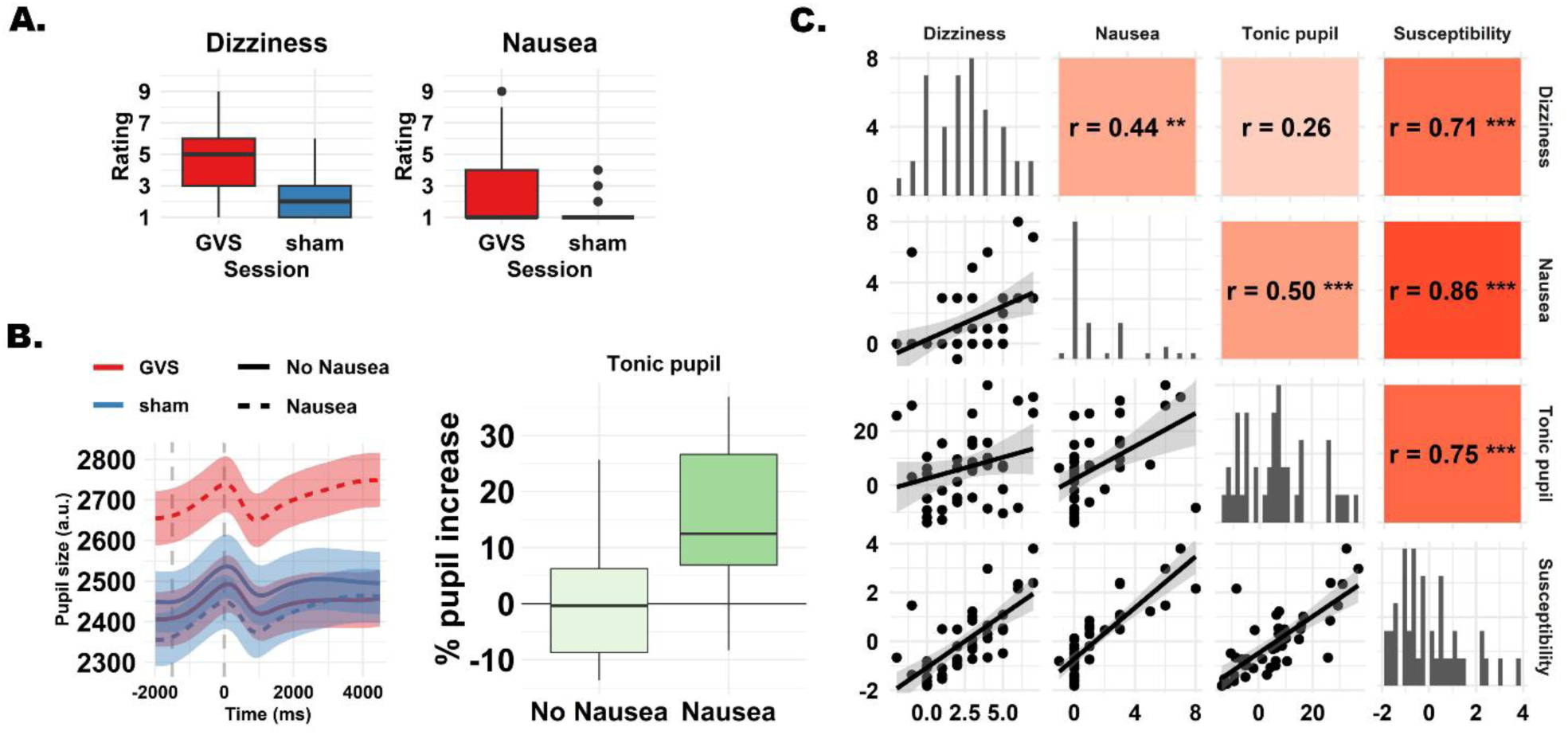
Individual susceptibility to vestibular stimulation. **A. Subjective ratings**. Participants (N= 42) had to report whether they experienced illusion of motion (left) or nausea (right) after both experimental conditions (GVS vs sham). Most participants reported increased dizziness and nausea during GVS, to variable degree. **B. Tonic pupil size.** During GVS, and throughout the entire session, the pupils were overall larger than during sham. The left plot depicts mean (SEM) pupil size throughout the entire duration of the trials. There was, however, substantial interindividual variability: pupil size betrayed whether participants experienced motion sickness and nausea. Among the participants who reported some degree of nausea (N= 18), pupil size increased on average by 14.4%, suggesting an overall heightened arousal state. **C. A continuous index of susceptibility to GVS.** Both subjective reports (dizziness and nausea) and objective physiological measures (tonic pupil size) were highly correlated. To account for interindividual differences in the effects of GVS, we submitted these variables to principal components analysis. The first principal component provides a continuous index of susceptibility to GVS and thus a proxy for the strength of interoceptive perturbations. K means clustering built on the same variables further delineated a cluster of participants (N= 15) subject to particularly strong effects (used for depictions in the next figures).

On the other hand, non-specific itching and tingling sensations induced by the electrical stimulation did not differ between conditions (*t*(41)= -1.88, *p*= 0.067); if anything, itching was more intense during the sham session, showing that the control condition worked as intended in inducing arousal originating from the physical sensations elicited by the electrodes. When participants were asked to rate the overall pain and discomfort caused by the procedure, however, the two conditions were closely matched (*t*(41)= -0.27, *p*= 0.79). Results are depicted in **Supplementary Figure 1**.

We then assessed whether participants rated the images as expected based on the normative data used to select them, involving arousal and valence. Negative images had on average low valence (M= 1.71, SD= 0.65) and high arousal scores (M= 6.47, SD= 0.84). Neutral images had on average medium valence scores (M= 5.09, SD= 0.74) and medium arousal scores (M= 3.45, SD= 0.68). Positive images had on average high valence (M= 6.94, SD= 1.08) and low arousal scores (M= 2.94, SD= 1.20). Note that we reframed the question exploring arousal in terms of elicited physiological arousal, so that positive images were the most calming and negative images were the most activating. Indeed, the three categories were all clearly separable in terms of both valence and arousal scores.

### GVS is associated with larger tonic pupil size

The overall (tonic) size of the pupil reflects slowly changing, rather stable physiological processes (Pelagatti et al., 2025). In the framework of the Adaptive Gain Theory (Aston-Jones & Cohen, 2005), tonic pupil size can be regarded as an indirect measure of Locus Coeruleus baseline activity and correlates with the overall behavioral mode (i.e., more or less focused on the task or the environment). That said, pupil size remains a low-dimensional readout of several interacting neural networks beyond the locus coeruleus (Joshi & Gold, 2020; Strauch et al., 2022).

For each participant, we computed the mean pupil size during the 500 ms preceding the delivery of electrical stimulation (from -2000 ms to -1500 ms). We then computed the percentage increase of tonic pupil size during GVS with the formula (pupilGVS – pupilSHAM)/pupilSHAM. Overall, the pupil was significantly larger during GVS (*t*(41)= 3.04, *p*= 0.004), increasing by 6.5% on average (CI95%= 2.1% –10.5%). However, the distribution was clearly bimodal. Partly, this could be due to counterbalancing. Long, boring tasks lead to the depletion of attentional resources and small pupils, which is probably why the tonic increase was numerically less pronounced in participants who received GVS after sham (CI95%=-1.6% – 9.9% vs 2.7% – 14.3%); however, there was no significant difference between the two groups (*t*(40)= -1.04, *p*= 0.3). Rather, tonic increase was likely driven by the subset of participants who felt stronger interoceptive effects, up to motion sickness and nausea (**Figure 2B**). Among the participants who reported some degree of nausea (N= 18), pupil size increased on average by 14.4% (CI95%= 8.2% – 20.7%), whereas the remaining participants presented no definite pattern (CI95%= -3.7% – 4.2%). The between-groups comparison was significant (*t*(40)= 3.92, *p*< 0.001). Thus, GVS is associated with increased tonic pupil size, suggesting an overall heightened arousal state. However, the magnitude of the increase is a function of the individual response to GVS and may be conditional to being aware of interoceptive perturbations such as nausea and motion sickness (**Figure 2B**).

### Quantifying susceptibility to GVS better captures interoceptive perturbations

We capitalized on a rather common, but fixed, stimulation intensity. Although this is a widespread solution, it also leaves open the possibility that GVS may have been delivered below (or above) the individual threshold. To account for this possibility, we therefore computed a measure of susceptibility to GVS. We used both the subjective ratings presented above (i.e. the magnitude of illusions of movement and motion sickness) and a more objective biomarker (i.e., the increase in tonic pupil size by GVS). We decided to merge subjective and objective measures because each has known limitations. For example, subjective reports may be prone to response bias and are constrained by one’s interoceptive sensitivity (that is, the ability to detect and report internal bodily states). On the other hand, physiological measures such as tonic pupil size may be somewhat noisy and highly dependent on external variables (e.g., counterbalancing, quality of the eye tracking data). Despite these limitations, the three variables showed consistently positive correlations (0.26< r < 0.50). Principal Component Analysis (PCA) identified one component accounting for 60% of their overall variance (PC1, eigenvalue= 1.344). This component loaded equally and positively onto the three constructs, and its scores thus provided a continuous index of susceptibility to GVS. We used this continuous measure for statistical modelling (**Figure 2C**). We additionally performed kmeans clustering on the original variables using k= 2. The procedure aims to delineate k groups with the constraint that differences must be maximized between groups and minimized within groups – i.e. groups must be homogeneous internally but also clearly separated. Two distinct clusters – one of GVS responders (N= 15, all experiencing nausea) and one of quasi-responders (N= 27, see below for evidence that GVS was effective in this group) – were identified along the continuum delineated by PC1. We used this solution to depict the results whenever the modelling pointed to individual susceptibility as an important modulator.

### The effects of GVS on basic oculomotor behavior are context-dependent

For all inferential statistics, we used linear mixed effects models (Baayen et al., 2008; Bates et al., 2015). It is notoriously challenging and hotly debated how to specify an optimal matrix of random effects, since the maximal models justified by the experimental design (Barr et al., 2013) tend to have convergence issues and should be kept parsimonious instead (Matuschek et al., 2017). Here we used the heuristics provided by the R package *buildmer* (Voeten, 2019), which identify the most complex possible regression model that will still converge. We used this approach to identify a suitable specification for random effects, up to a maximal model which included: a random intercept per participant; random slopes for Category (Negative, Positive, Neutral), Session (GVS, sham), and Stimulation (online, offline); all interactions (up to three-way) for the random slopes. The fixed effects were then tested via the package *afex* (Singmann et al., 2023), which uses Satterthwaite’s approximation as the default to calculate p values. We refer to the supplementary materials for the full details of model specification as well as the full results.

We started by assessing whether basic exploration features, notably the average number of fixations per image (**Supplementary Figure 3**) or their mean duration (**Supplementary Figure 4**), differed across some of our experimental variables, but it was not the case. Total dwell time did not differ across categories, sessions, or stimulation conditions either (**Supplementary Figure 2**). Thus, these features were consistent across images throughout the entire experiment, being largely unaffected by GVS even in the most susceptible participants.

We therefore moved to the spatial features of the fixations, beginning with their center of gravity (x and y coordinates). In the horizontal plane there was a strong interaction between Session and Stimulation (*F*(1, 74761.27)= 11.41, *p*< 0.001) so that active GVS led to rightward displacements whereas active sham led to leftward ones (**Figure 3**). This might seem at odds with classical GVS findings, where a shift of attention towards the anode (left) is usually expected (Ferrè et al., 2013). However, here we measured oculomotor behavior, rather than a covert shift in spatial attention, and previous reports have documented increased nystagmus (fast phase) towards the cathodal side of the stimulation (Nguyen et al., 2022). This effect was not modulated by Susceptibility to GVS (*F*< 0.2); if anything, the effect was *reduced* in highly susceptible participants (**Figure 3B**). This reflexive shift is rather expected considering the GVS settings, and the absence of a modulation by Susceptibility suggests that effects are somewhat comparable even in the group of participants who did not explicitly report vestibular effects. In short, this finding reflects the well-known automatic oculomotor response to the electric stimulation of the vestibular system.

**Figure 3.**
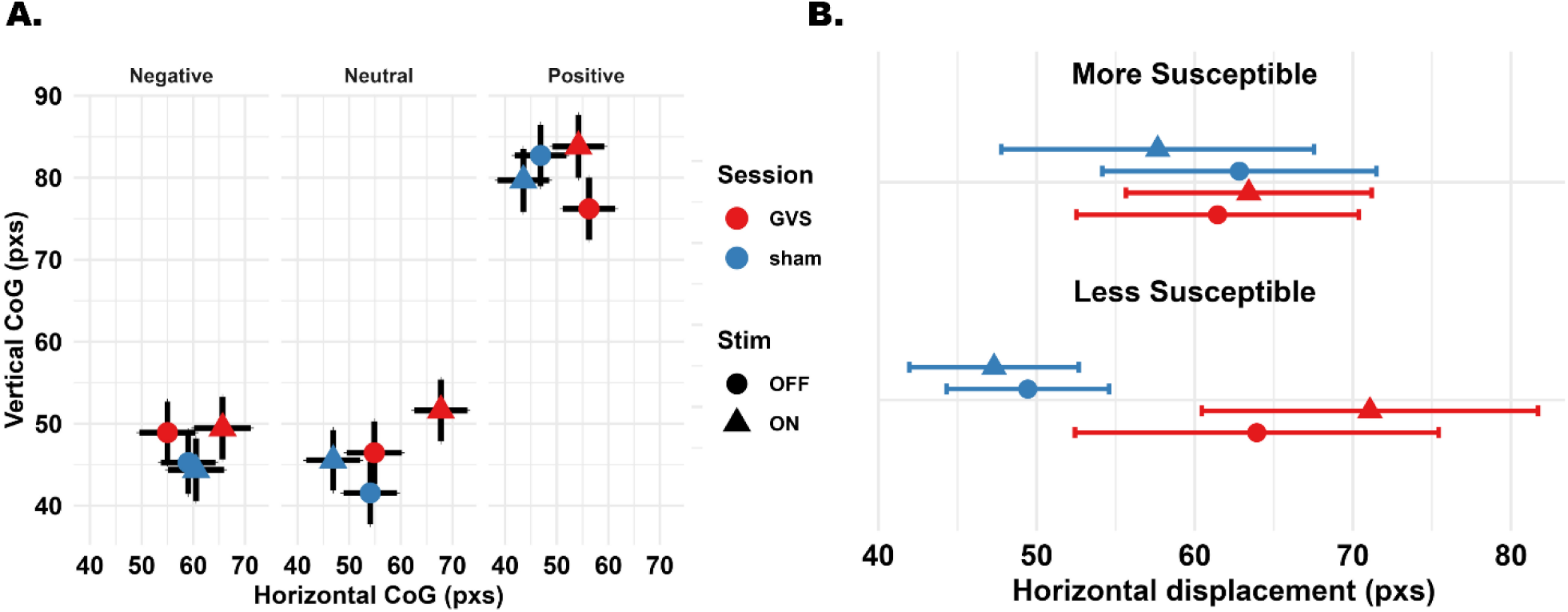
Effects of GVS on exploration spatial biases. **A. Fixation Center of Gravity (Mean ± SEM)**. GVS leads to a rightward displacement of fixations on the horizontal plane compared to sham, and there was also a small trend for GVS to induce upward biases. However, these effects were unequal across the three categories (also see Supplementary Figure 5). **B. Horizontal displacement as a function of susceptibility to GVS.** Mean (SEM) horizontal center of gravity for GVS and sham sessions, collapsed across image category. On average, GVS led to rightward displacement of the fixations’ center of gravity. There was no significant modulation of this bias by susceptibility to GVS, despite the visual trend.

However, this response was further modulated, to some extent, by Category (*F*(2, 74755.20)= 2.83, *p*= 0.059). Indeed, the interaction given by Stimulation and Session only holds within neutral images (*β*= -19.63, SE= 5.29, z= -3.71, *p*< 0.001), but not within negative (*β*= -9.15, SE= 5.27, z= -1.74, *p*= 0.082) or positive images (*β*= -2.06, SE= 5.26, z= -0.39, *p*= 0.696). In the vertical plane the evidence was less conclusive. The interaction between Session and Stimulation (*F*(1, 69379.43)= 3.65, *p*= 0.056) suggested a trend for online GVS to induce small upward displacements; however, even in this case the trend appeared to differ across categories (*F*(2, 67413.53)= 2.92, *p*= 0.054), since it was only significant within positive images (*β*= -11.5, SE= 3.79, z= -3.035, *p*= 0.002). There were no modulating effects of susceptibility to GVS (*F*< 0.1). Note that different categories are composed of different images and thus may present slightly different baseline centers of gravity (see **Supplementary Figure 5** for an alternative graphical depiction). Overall, GVS may induce subtle biases in visual exploration which likely depend on the stimulation parameters. However, these biases do not seem to vary as a function of the individual response to GVS. More importantly, the biases do not distribute homogeneously across all categories of images, showing that visual scanning is not completely overruled by non-specific, reflexive shifts in the position of the eyes, and that other processes continue contributing substantially to visual exploration.

### GVS alters reliance on perceptual salience

GVS affected the fixations’ center of gravity but did not do so similarly across all the categories of images. We therefore wondered whether one possible reason, besides the systematic shift of the eyes, was a different degree of reliance on bottom-up vs top-down mechanisms, which we probed in terms of average visual salience per fixation (Reynaud et al., 2021) (**Figure 4A**). There were two possibilities: 1) vestibular perturbations may increase external attention in an attempt to compensate for movement illusions, by “grabbing” onto visual features that are more stable and informative; 2) vestibular perturbations may increase instead attention towards internal processes, decoupling visual exploration from salient physical features (that is, a decreased average saliency per fixation). However, there was no main effect of Session (*F*(1, 37.56)= 0.26, *p*= 0.61). Interestingly, there was a non-significant trend for Susceptibility to GVS (*F*(1, 39.95)= 3.05, *p*= 0.088): on average (that is, across all sessions, categories, and stimulation conditions), highly susceptible participants tended to fixate more salient locations. Note that “susceptibility to GVS” was established based on subjective reports and pupil size, and not at all by features obtained from free viewing. The individual response to GVS is likely to vary primarily, albeit not solely, due to anatomical constraints. This trend, however, raises the fascinating speculation that the default pattern of visual exploration may also play a role, or perhaps betray a more pronounced visual dominance with respect to other senses. For example, participants who rely more heavily on informative visual cues may do so because of less stable and precise multisensory processes, which in turn would increase their susceptibility to GVS.

**Figure 4.**
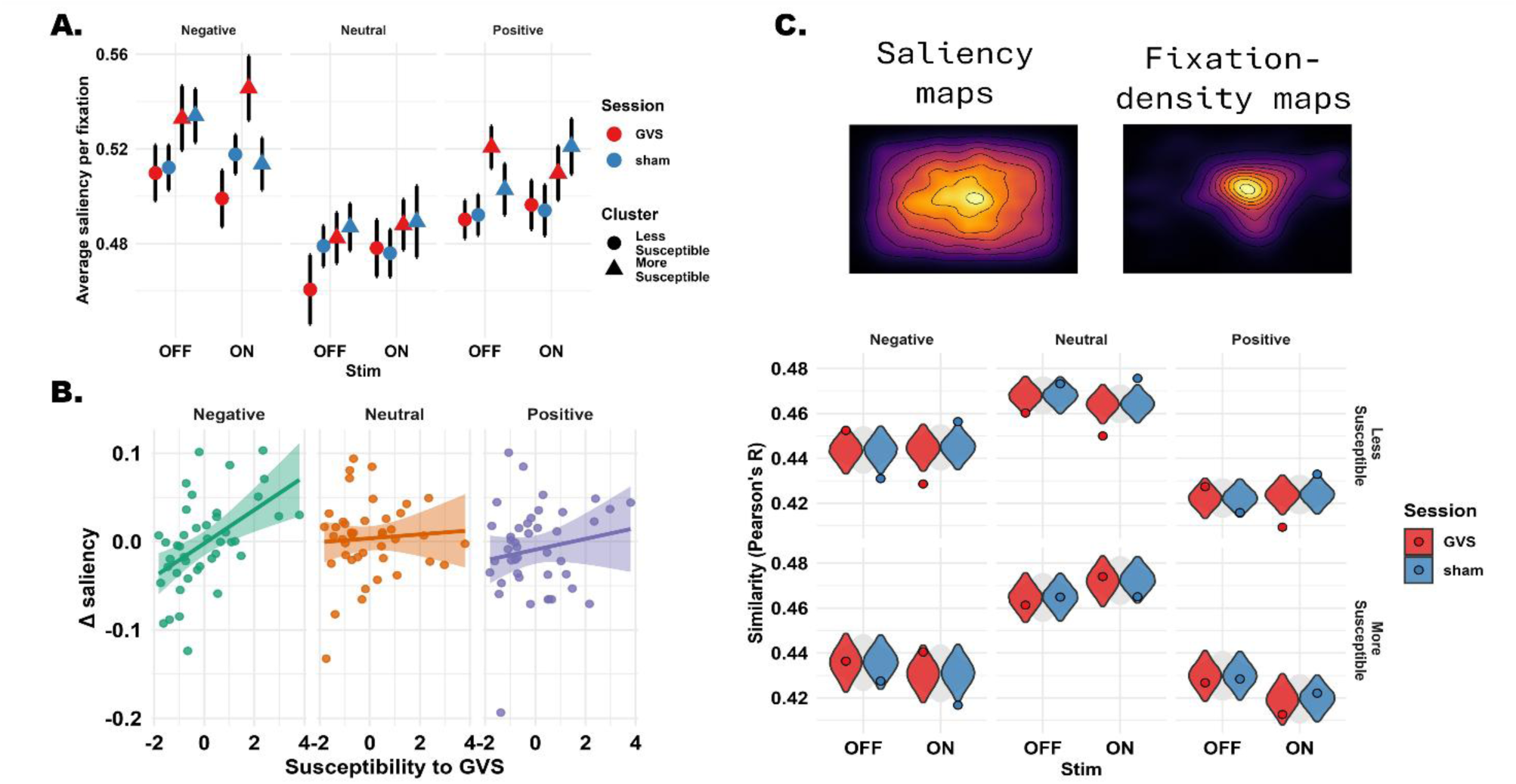
GVS alters reliance on low-level saliency. **A. Average saliency per fixation**. Mean (SEM) saliency per fixation across image categories, session, stimulation, and cluster of participants. We found a three-way interaction (Session x Category x Stimulation) which was, however, further qualified by the degree of susceptibility to GVS. Online GVS increased, for negative images only, the saliency associated with fixations, while online sham exerted the opposite effect; this effect was conditional to high values of susceptibility to GVS (here depicted a cluster of N= 15) and was reversed for less susceptible participants. **B. Correlation between susceptibility to GVS and** Δ **saliency.** We scored Δ saliency as the relative increase/decrease in the mean saliency per fixation from offline to online GVS, reflecting the different reliance on low-level properties of the images. Δ saliency was positively correlated with susceptibility to GVS when considering negative images and, marginally, positive images, but not neutral ones. The higher susceptibility to GVS, the higher the interoceptive perturbation, the stronger the reliance on low-level visual features in these conditions. **C. Comparison between fixation density maps and saliency maps.** The fixation data of all participants were collapsed (by the relevant conditions) to obtain more reliable fixation density maps (FDMs). FDMs were then compared directly with the theoretical saliency maps using Pearson’s R as measure of similarity. The value of R is depicted as a point in the plot. The interval depicted as a violin indicates the 95% confidence interval of the null model obtained via permutation test, in which we randomly shuffled the labels “GVS” or “sham” for each fixation within an image. Points falling outside this interval thus indicate R values that would be very extreme (<2.5%) under the null model. Online GVS consistently decreased reliance on low-level saliency in less susceptible participants. In highly susceptible participants, however, this trend was abolished, and even selectively reversed for negative images.

While no effects of Session were observed, and similarly to the analysis of spatial location presented above, there was a three-way interaction (Session x Category x Stimulation) which was, however, further qualified by the degree of susceptibility (*F*(2, 74664.79)= 3.88, *p*= 0.021). Post-hoc tests showed that, for negative images, online GVS increased the saliency associated with fixations while online sham exerted the opposite effect (*β*= -0.031, SE= 0.01, z= -3.17, *p*= 0.0015) respective to their offline conditions; this two-way interaction was not significant for neutral (*p*= 0.336) or positive images (*p*= 0.482). To further simplify the results, we repeated the analyses separately for each cluster of participants. The three-way interaction was not significant in the subgroup of participants who were less susceptible to GVS (*F*(2, 47166.71)= 0.17, *p*= 0.845). On the other hand, highly susceptible participants presented a significant three-way interaction (*F*(2, 27512.39)= 8.24, *p*< 0.001).

Another way to look at this complex interaction is to test whether Susceptibility to GVS and saliency show the same correlation strength across conditions. We therefore computed Pearson’s correlation coefficient between Susceptibility and the median saliency within each experimental cell. The only significant interaction was for negative images during online GVS (r= 0.451, *t*(40)= 3.19, *p*= 0.003), suggesting that participants experiencing stronger effect of GVS explored more salient low level visual features in this condition. There was also a trend, albeit not significant, for Susceptibility and salience to correlate within positive images during online GVS (r= 0.29, *t*(40)= 1.92, *p*= 0.062). For the remaining conditions, correlations were consistently positive but low (all *ps* >0.2). Because there is variability across images and categories with respect to the mean saliency per fixation, and to remain more faithful to the results above (which suggest an interaction between Session and Stimulation for negative images only) we scored Δ saliency as the relative increase/decrease from offline to online GVS stimulation (ON – OFF, **Figure 4B**). This reframing confirmed that Δ saliency increased significantly during online GVS selectively for negative images (r= 0.501, *t*(40)= 3.71, *p*< 0.001); the correlation was significantly larger than that observed for neutral images (z= 2.41, *p*= 0.016) and, marginally, for positive images (z= 1.96, *p*= 0.05).

Finally, we assessed the overall similarity (Pearson’s correlations) between fixation-density maps (FDM) and saliency maps (**Figure 4C**). We merged all participants within a cluster (more or less susceptible to GVS) to obtain robust FDRs; we then computed their similarity with the respective saliency maps (thus depending on each image), separately for each Session and Stimulation condition; we finally averaged the results of different images by Category, Session, and Condition. We set up a permutation test to probe statistical significance: we randomly shuffled for 50k times the labels “GVS” or “sham” separately for each image, Cluster, and Stimulation condition (ON vs OFF), then repeated the procedure above to create the distribution of null models depicted in **Figure 4C**. Results were highly consistent with the analysis concerning the average saliency per fixation. Online GVS consistently decreased the mean saliency per fixation in the group of participants who were less susceptible to GVS (all *ps*<0.001). On the other hand, highly susceptible participants tended to present the opposite pattern which, for negative images only, approached significance (*p*= 0.069); in their case, online sham led to a decreased reliance on visual saliency instead (*p*= 0.013). Finally, we scored the interaction contrast for Session x Stim x Cluster in our permutated null models and compared it with the original test score, separately for each image Category (corresponding to **Figure 4B** but also including the difference in slope between GVS and sham). Confirming previous analyses, the interaction was significant for negative (*β*= 0.064, *p*= 0.003), but not positive (*β*= 0.027, *p*= 0.085) or neutral images (*β*= 0.025, *p*= 0.098).

To summarize, the results concerning the spatial distribution of fixations were not entirely consistent with a purely reflexive, constant bias induced by GVS. Here we show that one important piece of the puzzle may be a different reliance on informative visual features while experiencing vestibular perturbations (see Castellotti et al., 2022, 2023), for evidence of gaze and attentional capture by information-optimal features). Online GVS generally *decreased*, in the less susceptible participants, the average saliency per fixation. This trend fits well with the systematic shift of the eyes described in the previous section: a constant bias that offsets fixations is likely to turn the eyes away from salient points, if those were the target of visual exploration. On the contrary, fixations landed on average on more salient and informative visual features in highly susceptible participants. Previous studies have shown increased visual capture by GVS in the context of multisensory illusions (Lopez, 2016; Lopez et al., 2010; Ponzo et al., 2018), which has been chiefly interpreted in terms of increased weighting of visual signals. Other groups found the exact opposite (Ferrè et al., 2015). Here we put forward the idea that – in addition to the differences in stimulation protocols – individual responses to GVS may play a critical role. Our results also raise the question as to whether visual capture may be associated with (or the consequence of) a qualitatively different pattern of visual exploration: favoring more informative features allows one a more thorough analysis of the visual scene, which in turn would offer, during vestibular perturbations, a more stable representation of the surrounding environment, particularly desirable for susceptible participants. Susceptible participants also presented increased tonic pupil size, as measured via an independent task. This, too, points towards a qualitatively different behavioral mode in highly susceptible participants (Aston-Jones & Cohen, 2005).

Finally, here we also show that this trend is context-sensitive, confirming previous claims that valence does play a role (Ponzo et al., 2018; Preuss et al., 2014). Visual exploration during online GVS was more thorough when images were highly salient (negative valence), further increasing a pre-existing bias towards low-level features and, concurrently, decreasing the role of endogenous processes. One previous study indeed found that caloric vestibular stimulation modulates mood and affective control (measured through an affective go/no-go task) during image viewing. A different exploration of the images, again, may at least partially account for these results. We therefore moved to assessing the phasic pupillary responses to GVS and, particularly, different image categories.

### GVS disrupts phasic pupillary responses to affective images

In contrast to tonic pupil size, *phasic* pupil responses are brief, event-locked constrictions/dilations reflecting transient locus coeruleus bursts that enhance processing of task-relevant stimuli when engaged. We start by assessing the effects of online stimulation – that is, using a baseline immediately prior to the electrical stimulation. We used linear mixed effects models for each timepoint, only using a random intercept for Participant to avoid convergence issues. We strive for clarity in this section: clusters are reported if significant at the *p*= 0.05 threshold unless otherwise stated; we refer the readers to **Supplementary Figure 6** for a complete overview of the effects tested and their statistics. We probed the fixed effects of Stimulation (ON vs OFF), Session (sham vs GVS) and Susceptibility to GVS. There was, unsurprisingly, a strong effect of the electrical stimulation (both sham and GVS) which, when delivered, caused the pupils to dilate after less than 400 ms and throughout the entire duration of the trial (-1120 ms – 4500 ms, fdr corrected). At the uncorrected level, however, multiple clusters of significant effects were also found. Most notably, Susceptibility interacted with both Stimulation and Session in the 1680-2680 ms and 2940-4500 ms windows. Chiefly, in the late stages of the trial pupil size dilated more for susceptible participants, and selectively so for online GVS, in keeping with their increased tonic pupil size (**Figure 5A**).

**Figure 5.**
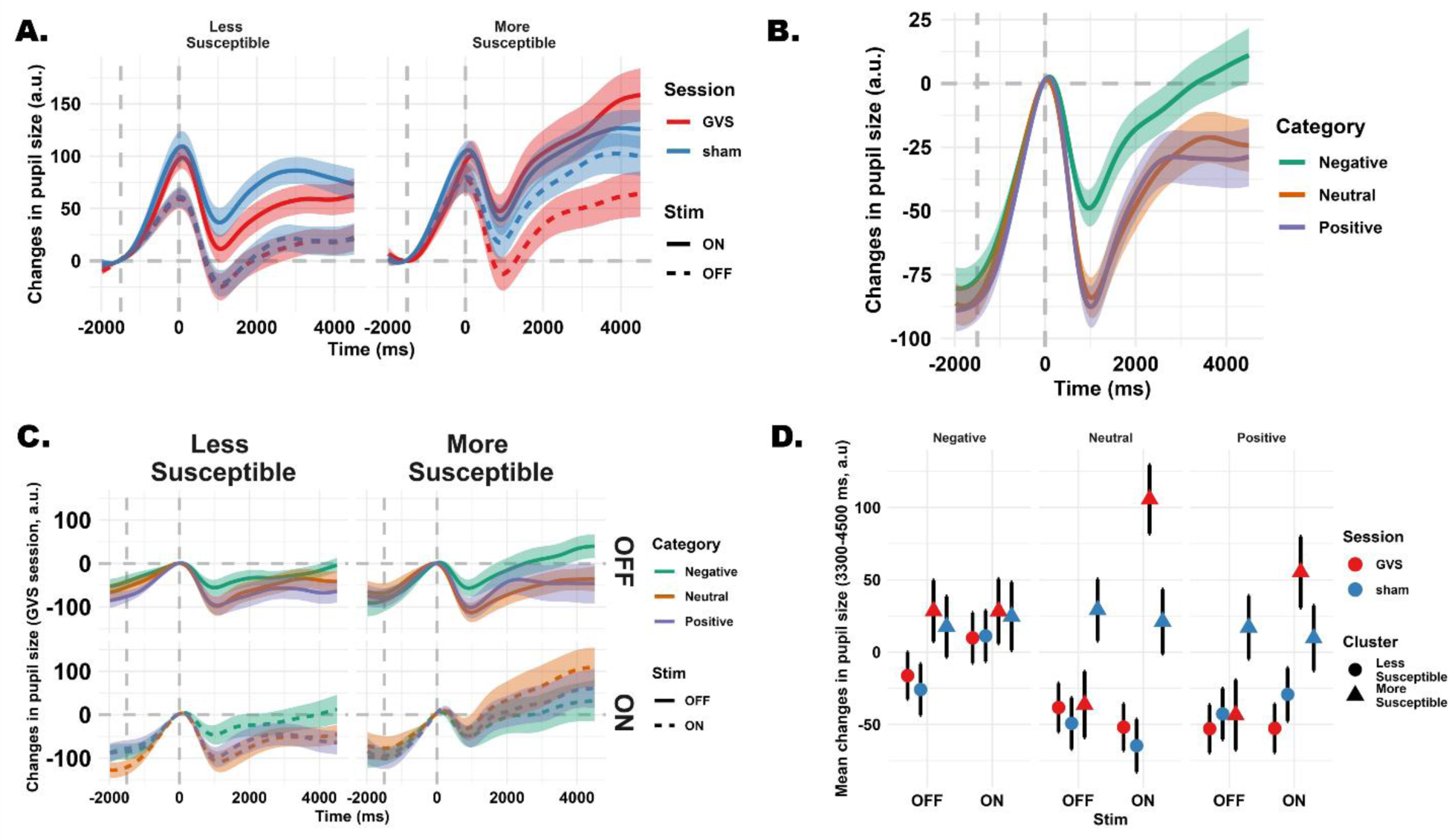
Phasic pupillary responses. **A. Change in pupil size as a function of time (mean ± SEM).** Electric stimulation (ON) induces a strong pupil dilation soon after 400ms; in the late stages of the trial, pupil size dilated more for susceptible participants during online GVS. **B. Overall affective pupillary responses.** Mean (SEM) changes in pupil size are depicted separately for the three image categories (collapsed across participants and conditions). Results show the classic pupil psychosensory response to the affect conveyed by the images, with negatively-valenced images associated with increased dilation. **C. Affective pupillary responses during vestibular stimulation.** Same as (B) but divided by cluster of participants (e.g., highly susceptible, N= 15) and stimulation (online vs offline); the plot only refers to pupil size during GVS (see Supplementary Figure 8 for sham). Affective pupil responses were generally preserved during sham and in less susceptible participants; responses were however abolished in highly susceptible participants during online GVS. **D. Mean pupil dilation (late psychosensory response).** The mean (SEM) dilation/constriction with respect to baseline is depicted across image categories, session, stimulation, and cluster of participants. Online GVS caused, in highly susceptible participants, the largest pupil dilation with respect to the offline condition, selectively for neutral images.

Since these windows are fully within the image presentation phase, we move to assessing phasic responses locked to the presentation of the images and we add the image Category as fixed factor (**Supplementary Figure 7**).

First, there was a clear effect of Category (**Figure 5B**). Most importantly for the current study, negative images were associated with larger dilations (or, rather, smaller constrictions) in two separate clusters, 400-2060 ms and 2600-4500 ms (fdr corrected). Results are consistent with the idea that emotional visual processing unfolds over multiple temporal stages, with early components reflecting rapid detection of salient information, and later components reflecting sustained elaboration (Olofsson et al., 2008). There were no differences between neutral and positive images in the context of this study. Because pupil size was unable to discriminate positive affect, we discarded this category of images to simplify and make more interpretable subsequent analyses.

At the fdr-corrected level, Susceptibility interacted with Session (940-1740 ms and 3200-4500 ms) and with both Session and Stimulation (320-4500 ms), chiefly reflecting the results highlighted above of increased phasic dilation for online GVS in highly susceptible participants. More importantly, there was an interaction (3300-4500 ms) between Susceptibility, Session, and Stimulation, for the specific contrast neutral vs negative images. In this window, the t-values ranged between 3.94 and 6.62 (0.01 > *p* > 0.047, **Figure 5C** and **Supplementary Figure 7**), though the interaction did not survive fdr correction across multiple timepoints. For a simpler interpretation, we ran the analyses separately for the two clusters of participants. In the group of participants that were less susceptible to GVS, there was only a main effect of Category (negative images being associated with larger pupil dilation) between 460 and 4500 ms (fdr corrected). For the most susceptible participants there was no main effect of Category but a three-way interaction between Category, Session, and Stimulation (300-4500 ms) suggesting that pupil dilation to negative images was selectively abolished during online GVS (**Figure 5C**, **Supplementary Figure 8**). A closer look at the data suggests that this may be the result of neutral images being associated with a particularly large pupil dilation during online GVS in highly susceptible participants. We scored the mean pupil size in the 3300-4500 window (**Figure 5D**) and submitted it to LMEMs analyses as detailed above (thus including random slopes that improved model fit, as guided by *buildmer*). There was a complex 4-way interaction for this scalar value (*F*(1, 11121.55)= 5.85, *p*= 0.016): only highly susceptible participants showed an interaction between Category, Stimulation, and Session (*β*= 157, SE= 53.71, z= -2.93, *p*= 0.0034). Further decomposition showed that only within neutral images online GVS caused the largest pupil dilation with respect to the offline condition (*β*= -142, SE= 37.9, z= -3.77, *p*< 0.001).

It is worth remembering that Susceptibility to GVS includes one prominent pupil feature: tonic pupil size. Susceptible participants were also characterized by larger baseline pupil size in the GVS session, which is known to affect the magnitude of phasic responses. On the other hand, larger pupils appear to be a defining feature of participants experiencing nausea, which is why tonic and phasic responses are ultimately intertwined and difficult to truly disentangle. That said, we note that results were unchanged when we regressed out from each timepoint and trial the respective baseline value (**Supplementary Figure 9**).

In the online condition, electrical currents were presented before image presentation. It is reasonable to expect larger, stimulation-related phasic dilation in the subset of participants who felt stronger GVS effects, as we did; it is less clear, however, why neutral images would be comparatively more affected. One possibility consists of potential ceiling effects for changes in pupil size: ceiling effects could prevent any further dilation caused by negative images (**Supplementary Figure 10**). Alternatively, the lack of affective context in the neutral condition may have allowed a more robust autonomic effect of GVS, whereas an affective context could have prompted a state of enhanced cognitive control. On the other hand, it is unlikely that altered pupillary responses are the consequence of a different scanning of the images (i.e., more driven by salience and low-level features) because images in this task were very tiny, precisely to avoid eye movements. Still, it is particularly striking the connection between this finding and that of increased reliance on low-level saliency in a separate free-viewing task. Both measures – visual saliency and phasic pupil responses – were selectively altered during online GVS and for negative images. There was a small correlation between ΔSaliency in the IVT and the gain in pupil size observed here for neutral images (**Supplementary Figure 11**). Vestibular stimulation (Preuss et al., 2014) has been found to modulate mood and affective processing before, including affective control. GVS has been associated with soothing and calming effects, potentially capable to reduce anxiety in healthy adults (Pasquier et al., 2019). Consistently, our results suggest that both tonic and phasic arousal-linked mechanisms may ultimately underlie these findings.

## Discussion

We administered GVS to 42 healthy participants freely exploring affective images, while recording their eye movements and pupil size in separate tasks. We demonstrate that vestibular perturbations reshape visual exploration beyond purely reflexive shifts of the eyes. This effect is strongly dependent on individual susceptibility to GVS and image content, and is accompanied by a distinctive disruption of pupillary dynamics.

First, GVS was effective in inducing variable degrees of dizziness and nausea. Participants reported, anecdotally, feeling “mildly drunk” or as “rocking on a boat”, further corroborating the interoceptive nature of this manipulation. As a group, they also showed a stark increase in tonic pupil size. However, there was substantial heterogeneity in both subjective reports and more objective biomarkers (pupil size): only a subset of participants presented clear illusions of movement, nausea, and larger pupil size, effectively driving the average of the entire sample. We argue that individual susceptibility is key to understanding the high heterogeneity of neurostimulation studies (Chew et al., 2015). The same holds for peripheral stimulation techniques: the effect of auricular vagal stimulation on pupil size, for example, has been debated (Pervaz et al., 2025), with effects that are known to be dose-dependent (Capone et al., 2021). The majority of studies rightfully lamented the heterogeneity of stimulation parameters and methodological choices (Pliego et al., 2025), whereas the topic of individual differences has been approached more reluctantly. Individual susceptibility was in our case, however, also associated with radically different behavioral patterns – as would be expected, in the framework of the adaptive gain theory (Aston-Jones & Cohen, 2005), in the presence of larger tonic pupil size. We indeed observed a remarkable dissociation involving the visual exploration of images, specifically the reliance on low-level features.

At the group level, GVS caused a sustained bias in the displacement of horizontal fixations that was consistent with the induced hemispheric asymmetries. However, this bias was unequal across the different categories, demonstrating its context-dependent nature. In particular, the expected shift only held within neutral images, pointing to affective valence as a key factor modulating visual exploration.

This systematic bias could be detected in participants who felt less interoceptive perturbations as well, showing that the vestibular system was stimulated as intended in all participants. This is important because it provides means to tease apart the effects of stimulating the vestibular system from the arguably qualitatively different condition in which strong interoceptive perturbations arise. Accordingly, this extrinsic (GVS-induced) bias disrupted the coupling between gaze and low-level visual features. With online GVS, and across all image categories, the fixations landed at less physically salient locations, a result consistent with the disruption of a normal egocentric reference frame (Fink et al., 2003). Translating this finding to multisensory contexts is not straightforward; however, we argue that this pattern of results is consistent with a reduced weight assigned to vision (Ferrè et al., 2015). There are two, non-mutually exclusive directions: fixations may land on less informative locations, thus lowering the impact of vision as consequence; a decreased reliance on vision may lead to fixations that are decoupled from physical saliency. We tend, however, to favor a scenario in which the two processes interact dynamically. Importantly, the cluster of participants who presented strong interoceptive sensations arising from GVS also presented the opposite behavioral pattern. This was the case despite 1) an identical intensity of the electrical stimulation and 2) individual susceptibility was defined based on the results of independent ratings and an independent task. Highly susceptible participants increased their reliance on salient and informative locations, especially for negative images, which is consistent with increased visual capture instead (Lopez et al., 2010; Ponzo et al., 2018). Strong interoceptive perturbations may cause participants to increase their reliance on salient visual cues and thus explore more thoroughly the visual scene, perhaps in attempt to leverage more stable features of the environment, in principle better suited to mitigate the dizziness and anchor themselves onto the external world. While methodological differences (i.e., the duration of stimulation) may still play a role, our findings can therefore reconcile nicely the different results in the literature concerning visual capture: decreased and increased visual capture can coexist depending on the individual response to GVS. In addition to the classic view of vestibular stimulations biasing eye movements “reactively”, moreover, our results put forward the novel idea that at least part of these changes may be “adaptive”. Arguably this can only be appreciated when moving beyond artificial and strictly controlled settings and visual stimuli to adopt more naturalistic, free viewing conditions of complex stimuli.

Larger tonic pupil size by itself is not necessarily sufficient to disentangle conflicting hypotheses on whether attention is directed toward internal or external states (Pelagatti et al., 2025). Both focusing on internal states – especially when accompanied by nausea and worry – and increased alertness to the environment can cause larger pupils. The use of saliency-based metrics in the present study allows us to disentangle the two accounts in favor of the latter. Saliency-based metrics, indeed, are typically interpreted in terms of dynamic balance between bottom-up and top-down processes; having observed increased reliance on saliency with increasing susceptibility suggests that environmental alertness is predominant at the expense of more evaluative, top-down aspects. This is also consistent with the finding that negatively-valenced images are prominently affected. Our results globally show that interoceptive perturbations driven by GVS dynamically reshape the balance between bottom-up and top-down attentional processes, with implications for affective processing and adaptive behavior. Interestingly, one study have shown that viewing emotionally negative images akin to ours can improve the discrimination of vestibular input (Preuss et al., 2015), suggesting that the link may be bidirectional.

We acknowledge that our study has several limitations in this regard. The image pool was limited to 12 stimuli per category, and there was substantial variability between images. The variability was particularly extreme for positive images. While this limitation was necessary in the present work, future studies should ensure that the stimuli generalize. Negative images, however, are more meaningfully affected by increased environmental alertness, given their potential for threat. Furthermore, negative images are more likely to be affected by the “affective gap” caused by ongoing evaluative (top-down) processes. The affective gap consists of a decoupling between gaze and visual saliency when the emotional context elicits top-down interference. The assessment of phasic changes in pupil size highlighted indeed the signature pupil dilation associated with emotional content (Bogdanova et al., 2022; Dureux et al., 2021; Gilzenrat et al., 2010; Murphy et al., 2011), and demonstrated that, in the present study, it was selective for negative images. Highly susceptible participants presented, during online GVS, an overall disrupted pattern of changes in pupil size in which, crucially, this signature dilation was abolished. We remain cautious in the interpretation of these results because ceiling effects may have a role: highly susceptible participants presented larger baseline pupil size and stronger phasic responses overall, which may have hindered subtle effects. Results are, however, consistent with previous studies showing the potential for vestibular stimulation techniques to alter mood and affective control in a valence-specific way (Preuss et al., 2014, 2015). GVS has been associated with soothing and calming effects, potentially capable of reducing anxiety in healthy adults (Pasquier et al., 2019) or even manic symptoms in bipolar patients (according to single-case reports Dodson, 2004). Our results suggest that both tonic and phasic arousal-linked mechanisms may accompany these effects. The disruption of the signature effect of negative images on phasic pupil responses, in particular, suggests that evaluative top-down processes sensitive to emotional content and behavioral relevance are hindered or diluted by an overall pupil dilation caused by the interoceptive perturbation, which may lead to enhanced cognitive control. The neurological bases of the functional links between vestibulo-autonomic control and affect (notably anxiety) have been described extensively: they may revolve around the parabrachial nucleus network and its reciprocal interactions with the amygdaloid nucleus, hypothalamus, and other limbic cortices more generally (Balaban, 2004; Balaban & Thayer, 2001). Additionally, GVS may influence locus coeruleus activity, which regulates arousal, neuromodulatory signaling (primarily noradrenaline), and adaptive gain to optimize behavioral performance (Aston-Jones & Cohen, 2005; Reynaud et al., 2021), similarly to what has been proposed for vagal stimulation (Sharon et al., 2021). More direct measures of neural activity will be necessary to substantiate the causal role of these circuits.

How, then, do we really look at the world when experiencing nausea? Perception is not a mere assembly of physical features. It is a creative act which merges low- and high-level components. Among the latter, top-down processes tied to emotional content color our phenomenological experience and alter the way information is sampled from the environment: affective predictions support and dynamically interact with vision early on (Barrett & Bar, 2009). Interoceptive perturbations largely override these naturally occurring processes and may dilute affective processing tied to specific visual stimuli, especially in participants that are particularly susceptible. The consequence is that the “colors” of visual experience are altered and, in a sense, they fade away. This further propagates into a qualitatively different visual scanning of salient features. While perception may become dull, on the other hand, it may also gain a better hold onto reality in its physical, objective counterpart. To summarize, our results unveil the continuous interplay between interoceptive and exteroceptive processing, and show that how we perceive the environment fundamentally resonates with how we feel.

Vestibular stimulation is increasingly used for rehabilitation, as well as in engineering to mitigate virtual reality sickness or to create more immersive gaming experiences (Cevette et al., 2014; Marchand et al., 2025). Here we show that, depending on the setting and individual susceptibility, its effects may be more profound than anticipated. We also show that GVS may be a powerful tool for basic research. It allows one to draw, with enhanced experimental control, causal inferences on the continuous dialogue between bodily processes, perception, and decision-making.

## Acknowledgements

The authors are grateful to Dr. Elena Tonolli and Dr. Gianpiero Monittola for technical assistance. The project has been funded by the European Union - NextGenerationEU, Investment line 1.2 “Funding projects presented by young researchers”, title "Behavioral, autonomic, and Electrophysiological SignaTures of Vestibular Stimulation (BEST-VS)", CUP: E73C25000210001.

## Data availability statement

The relevant computer code as well as the raw data are available without restrictions at the following link: https://osf.io/dwjzr. The study was not formally preregistered.

## Author Contributions

EB conceived the research question and secured funding. EB and IP designed the experiments. EB and IP collected, analyzed, and depicted the data. EB wrote the first draft with substantial input and revisions from IP and FHB.

## References

Angelaki, D. E., Klier, E. M., & Snyder, L. H. (2009). A vestibular sensation: Probabilistic approaches to spatial perception. Neuron, 64(4), 448–461. 10.1016/j.neuron.2009.11.010

Aston-Jones, G., & Cohen, J. D. (2005). AN INTEGRATIVE THEORY OF LOCUS COERULEUS-NOREPINEPHRINE FUNCTION: Adaptive Gain and Optimal Performance. Annual Review of Neuroscience, 28(1), 403–450. 10.1146/annurev.neuro.28.061604.135709

Baayen, R. H., Davidson, D. J., & Bates, D. M. (2008). Mixed-effects modeling with crossed random effects for subjects and items. Journal of Memory and Language, 59(4), 390–412. 10.1016/j.jml.2007.12.005

Balaban, C. D. (2004). Projections from the parabrachial nucleus to the vestibular nuclei: Potential substrates for autonomic and limbic influences on vestibular responses. Brain Research, 996(1), 126–137. 10.1016/j.brainres.2003.10.026

Balaban, C. D., & Thayer, J. F. (2001). Neurological bases for balance–anxiety links. *Journal of Anxiety Disorders*, The Interface of Balance Disorders and Anxiety, 15(1), 53–79. 10.1016/S0887-6185(00)00042-6

Barr, D. J., Levy, R., Scheepers, C., & Tily, H. J. (2013). Random effects structure for confirmatory hypothesis testing: Keep it maximal. Journal of Memory and Language, 68(3), 255–278. 10.1016/j.jml.2012.11.001

Barrett, L. F., & Bar, M. (2009). See it with feeling: Affective predictions during object perception. Philosophical Transactions of the Royal Society B: Biological Sciences, 364(1521), 1325–1334. 10.1098/rstb.2008.0312

Barrett, L. F., & Simmons, W. K. (2015). Interoceptive predictions in the brain. Nature Reviews Neuroscience, 16(7), 419–429. 10.1038/nrn3950

Bates, D., Mächler, M., Bolker, B., & Walker, S. (2015). Fitting Linear Mixed-Effects Models Using lme4. Journal of Statistical Software, 67(1), 1–48. 10.18637/jss.v067.i01

Binda, P., & Gamlin, P. D. (2017). Renewed Attention on the Pupil Light Reflex. Trends in Neurosciences, 40(8), 455–457. 10.1016/j.tins.2017.06.007

Bisiach, E., Rusconi, M. L., & Vallar, G. (1991). Remission of somatoparaphrenic delusion through vestibular stimulation. Neuropsychologia, 29(10), 1029–1031. 10.1016/0028-3932(91)90066-H

Blini, E., Arrighi, R., & Anobile, G. (2024). Pupillary manifolds: Uncovering the latent geometrical structures behind phasic changes in pupil size. Scientific Reports, 14(1), 27306. 10.1038/s41598-024-78772-x

Blini, E., Tilikete, C., Chelazzi, L., Farnè, A., & Hadj-Bouziane, F. (2020). The role of the vestibular system in value attribution to positive and negative reinforcers. Cortex, 133, 215–235. 10.1016/j.cortex.2020.09.004

Blini, E., Tilikete, C., Farnè, A., & Hadj-Bouziane, F. (2018). Probing the role of the vestibular system in motivation and reward-based attention. Cortex, 103, 82–99. 10.1016/j.cortex.2018.02.009

Blini, E., & Zorzi, M. (2023). Pupil size as a robust marker of attentional bias toward nicotine-related stimuli in smokers. Psychonomic Bulletin & Review, 30(2), 596–607. 10.3758/s13423-022-02192-z

Bogdanova, O. V., Bogdanov, V. B., Miller, L. E., & Hadj-Bouziane, F. (2022). Simulated proximity enhances perceptual and physiological responses to emotional facial expressions. Scientific Reports, 12(1), Article 1. 10.1038/s41598-021-03587-z

Botvinick, M., & Cohen, J. (1998). Rubber hands ‘feel’ touch that eyes see. Nature, 391(6669), 756–756. 10.1038/35784

Capone, F., Motolese, F., Di Zazzo, A., Antonini, M., Magliozzi, A., Rossi, M., Marano, M., Pilato, F., Musumeci, G., Coassin, M., & Di Lazzaro, V. (2021). The effects of transcutaneous auricular vagal nerve stimulation on pupil size. Clinical Neurophysiology, 132(8), 1859–1865. 10.1016/j.clinph.2021.05.014

Cappa, S., Sterzi, R., Vallar, G., & Bisiach, E. (1987). Remission of hemineglect and anosognosia during vestibular stimulation. Neuropsychologia, 25(5), 775–782. 10.1016/0028-3932(87)90115-1

Carmona, J. E., Holland, A. K., & Harrison, D. W. (2009). Extending the functional cerebral systems theory of emotion to the vestibular modality: A systematic and integrative approach. Psychological Bulletin, 135(2), 286–302. 10.1037/a0014825

Castellotti, S., Castaldi, E., Blini, E., & Arrighi, R. (2025). Pupil size as a biomarker of cognitive (dys)functions: Toward a physiologically informed screening of mental states. Acta Psychologica, 253, 104720. 10.1016/j.actpsy.2025.104720

Castellotti, S., Montagnini, A., & Del Viva, M. M. (2022). Information-optimal local features automatically attract covert and overt attention. Scientific Reports, 12(1), 9994. 10.1038/s41598-022-14262-2

Castellotti, S., Petrizzo, I., Arrighi, R., & Blini, E. (2025). Dimensionality reduction techniques in pupillometry research: A primer for behavioral scientists. Behavior Research Methods, 57(12), 337. 10.3758/s13428-025-02786-0

Castellotti, S., Szinte, M., Del Viva, M. M., & Montagnini, A. (2023). Saccadic trajectories deviate toward or away from optimally informative visual features. iScience, 26(8), 107282. 10.1016/j.isci.2023.107282

Ceunen, E., Vlaeyen, J. W. S., & Van Diest, I. (2016). On the Origin of Interoception. Frontiers in Psychology, 7, 743. 10.3389/fpsyg.2016.00743

Cevette, M., Stepanek, J., & Galea, A. (2014). Galvanic vestibular stimulation system and method of use for simulation, directional cueing, and alleviating motion-related sickness (United States Patent No. US8718796B2). https://patents.google.com/patent/US8718796B2/en

Chew, T., Ho, K.-A., & Loo, C. K. (2015). Inter- and Intra-individual Variability in Response to Transcranial Direct Current Stimulation (tDCS) at Varying Current Intensities. Brain Stimulation, 8(6), 1130–1137. 10.1016/j.brs.2015.07.031

Cinetto, S., Blini, E., Zangrossi, A., Corbetta, M., & Zorzi, M. (2025). Spatial regularities in a closed-loop audiovisual search task bias subsequent free-viewing behavior. Psychonomic Bulletin & Review, 32(6), 2977–2989. 10.3758/s13423-025-02703-8

Craig, A. D. (2002). How do you feel? Interoception: the sense of the physiological condition of the body. Nature Reviews Neuroscience, 3(8), 655–666. 10.1038/nrn894

Critchley, H. D., & Garfinkel, S. N. (2017). Interoception and emotion. *Current Opinion in Psychology*, Emotion, 17, 7–14. 10.1016/j.copsyc.2017.04.020

Deroualle, D., & Lopez, C. (2014). Toward a vestibular contribution to social cognition. Frontiers in Integrative Neuroscience, 8, 16. 10.3389/fnint.2014.00016

Dodson, M. J. (2004). Vestibular stimulation in mania: A case report. *Journal of Neurology*, Neurosurgery & Psychiatry, 75(1), 168–169.

Dureux, A., Blini, E., Grandi, L. C., Bogdanova, O., Desoche, C., Farnè, A., & Hadj-Bouziane, F. (2021). Close facial emotions enhance physiological responses and facilitate perceptual discrimination. Cortex, 138, 40–58. 10.1016/j.cortex.2021.01.014

Engbert, R., & Kliegl, R. (2003). Microsaccades uncover the orientation of covert attention. Vision Research, 43(9), 1035–1045. 10.1016/s0042-6989(03)00084-1

Ferrè, E. R., Berlot, E., & Haggard, P. (2015). Vestibular contributions to a right-hemisphere network for bodily awareness: Combining galvanic vestibular stimulation and the “Rubber Hand Illusion.” Neuropsychologia, 69, 140–147. 10.1016/j.neuropsychologia.2015.01.032

Ferrè, E. R., & Haggard, P. (2016). The vestibular body: Vestibular contributions to bodily representations. Cognitive Neuropsychology, 33(1–2), 67–81. 10.1080/02643294.2016.1168390

Ferrè, E. R., & Haggard, P. (2020). Vestibular cognition: State-of-the-art and future directions. Cognitive Neuropsychology, 37(7–8), 413–420. 10.1080/02643294.2020.1736018

Ferrè, E. R., Longo, M., Fiori, F., & Haggard, P. (2013). Vestibular modulation of spatial perception. Frontiers in Human Neuroscience, 7:, 660. 10.3389/fnhum.2013.00660

Fink, G. R., Marshall, J. C., Weiss, P. H., Stephan, T., Grefkes, C., Shah, N. J., Zilles, K., & Dieterich, M. (2003). Performing allocentric visuospatial judgments with induced distortion of the egocentric reference frame: An fMRI study with clinical implications. NeuroImage, 20(3), 1505–1517. 10.1016/j.neuroimage.2003.07.006

Finke, J. B., Roesmann, K., Stalder, T., & Klucken, T. (2021). Pupil dilation as an index of Pavlovian conditioning. A systematic review and meta-analysis. Neuroscience & Biobehavioral Reviews, 130, 351–368. 10.1016/j.neubiorev.2021.09.005

Fitzpatrick, R. C., & Day, B. L. (2004). Probing the human vestibular system with galvanic stimulation. Journal of Applied Physiology (Bethesda, Md.: *1985)*, 96(6), 2301–2316. 10.1152/japplphysiol.00008.2004

Gilzenrat, M. S., Nieuwenhuis, S., Jepma, M., & Cohen, J. D. (2010). Pupil diameter tracks changes in control state predicted by the adaptive gain theory of locus coeruleus function. *Cognitive, Affective*, & Behavioral Neuroscience, 10(2), 252–269. 10.3758/CABN.10.2.252

Hammam, E., Dawood, T., & Macefield, V. G. (2012). Low-frequency galvanic vestibular stimulation evokes two peaks of modulation in skin sympathetic nerve activity. Experimental Brain Research, 219(4), 441–446. 10.1007/s00221-012-3090-z

Harel, J., Koch, C., & Perona, P. (2007). Graph-Based Visual Saliency. In B. Schölkopf, J. Platt, & T. Hofmann (Eds.), Advances in Neural Information Processing Systems 19 (NIPS 2006) (No. 19; Issue 19, pp. 545–552). MIT Press. 20th Conference on Advances in Neural Information Processing Systems (NIPS). https://resolver.caltech.edu/CaltechAUTHORS:20160315-111145907

Hoover, A. E. N., & Harris, L. R. (2015). Disrupting Vestibular Activity Disrupts Body Ownership. Multisensory Research, 28(5–6), 581–590. 10.1163/22134808-00002472

Itti, L., & Koch, C. (2000). A saliency-based search mechanism for overt and covert shifts of visual attention. Vision Research, 40(10), 1489–1506. 10.1016/S0042-6989(99)00163-7

Joshi, S., & Gold, J. I. (2020). Pupil Size as a Window on Neural Substrates of Cognition. Trends in Cognitive Sciences, 24(6), 466–480. 10.1016/j.tics.2020.03.005

Karnath, H. O., Fetter, M., & Dichgans, J. (1996). Ocular exploration of space as a function of neck proprioceptive and vestibular input—Observations in normal subjects and patients with spatial neglect after parietal lesions. Experimental Brain Research. Experimentelle Hirnforschung. Expérimentation Cérébrale, 109(2), 333–342.

Kaspar, K., & König, P. (2011a). Overt Attention and Context Factors: The Impact of Repeated Presentations, Image Type, and Individual Motivation. PLOS ONE, 6(7), e21719. 10.1371/journal.pone.0021719

Kaspar, K., & König, P. (2011b). Viewing behavior and the impact of low-level image properties across repeated presentations of complex scenes. Journal of Vision, 11(13). 10.1167/11.13.26

Kurdi, B., Lozano, S., & Banaji, M. R. (2017). Introducing the Open Affective Standardized Image Set (OASIS). Behavior Research Methods, 49(2), 457–470. 10.3758/s13428-016-0715-3

Kwan, A., Forbes, P. A., Mitchell, D. E., Blouin, J.-S., & Cullen, K. E. (2019). Neural substrates, dynamics and thresholds of galvanic vestibular stimulation in the behaving primate. Nature Communications, 10(1), 1904. 10.1038/s41467-019-09738-1

Le Meur, O., & Baccino, T. (2013). Methods for comparing scanpaths and saliency maps: Strengths and weaknesses. Behavior Research Methods, 45(1), 251–266. 10.3758/s13428-012-0226-9

Lenggenhager, B., & Lopez, C. (2015). Vestibular contributions to the sense of body, self, and others. 10.25358/openscience-68

Lenggenhager, B., Lopez, C., & Blanke, O. (2007). Influence of galvanic vestibular stimulation on egocentric and object-based mental transformations. Experimental Brain Research, 184(2), 211–221. 10.1007/s00221-007-1095-9

Lenth, R. (2019). Emmeans: Estimated Marginal Means, aka Least-Squares Means. R package version 1.*4*.1.

https://CRAN.R-project.org/package=emmeans

Lopez, C. (2015). Making Sense of the Body: The Role of Vestibular Signals. Multisensory Research, 28(5–6), 525–557. 10.1163/22134808-00002490

Lopez, C. (2016). The vestibular system: Balancing more than just the body. Current Opinion in Neurology, 29(1), 74–83. 10.1097/WCO.0000000000000286

Lopez, C., Blanke, O., & Mast, F. W. (2012). The human vestibular cortex revealed by coordinate-based activation likelihood estimation meta-analysis. Neuroscience, 212, 159–179. 10.1016/j.neuroscience.2012.03.028

Lopez, C., Lenggenhager, B., & Blanke, O. (2010). How vestibular stimulation interacts with illusory hand ownership. Consciousness and Cognition, 19(1), 33–47. 10.1016/j.concog.2009.12.003

Malsburg, T. von der. (2019). *saccades: Detection of Fixations in Eye-Tracking Data*. https://github.com/tmalsburg/saccades

Marchand, S., Langlade, A., Legois, Q., & Séverac Cauquil, A. (2025). A wide-ranging review of galvanic vestibular stimulation: From its genesis to basic science and clinical applications. Experimental Brain Research, 243(5), 131. 10.1007/s00221-025-07079-8

Mast, F. W., Preuss, N., Hartmann, M., & Grabherr, L. (2014). Spatial cognition, body representation and affective processes: The role of vestibular information beyond ocular reflexes and control of posture. Frontiers in Integrative Neuroscience, 8, 44. 10.3389/fnint.2014.00044

Mathôt, S., Schreij, D., & Theeuwes, J. (2012). OpenSesame: An open-source, graphical experiment builder for the social sciences. Behavior Research Methods, 44(2), 314–324. 10.3758/s13428-011-0168-7

Matuschek, H., Kliegl, R., Vasishth, S., Baayen, H., & Bates, D. (2017). Balancing Type I error and power in linear mixed models. Journal of Memory and Language, 94, 305–315. 10.1016/j.jml.2017.01.001

Murphy, P. R., Robertson, I. H., Balsters, J. H., & O’connell, R. G. (2011). Pupillometry and P3 index the locus coeruleus–noradrenergic arousal function in humans. Psychophysiology, 48(11), 1532–1543. 10.1111/j.1469-8986.2011.01226.x

Nguyen, T. T., Kang, J.-J., & Oh, S.-Y. (2022). Thresholds for vestibular and cutaneous perception and oculomotor response induced by galvanic vestibular stimulation. Frontiers in Neurology, 13. 10.3389/fneur.2022.955088

Niu, Y., Todd, R., & Anderson, A. K. (2012). Affective salience can reverse the effects of stimulus-driven salience on eye movements in complex scenes. Frontiers in Psychology, 3. 10.3389/fpsyg.2012.00336

Oldfield, R. C. (1971). The assessment and analysis of handedness: The Edinburgh inventory. Neuropsychologia, 9(1), 97–113. 10.1016/0028-3932(71)90067-4

Olofsson, J. K., Nordin, S., Sequeira, H., & Polich, J. (2008). Affective picture processing: An integrative review of ERP findings. Biological Psychology, 77(3), 247–265. 10.1016/j.biopsycho.2007.11.006

Pasquier, F., Denise, P., Gauthier, A., Bessot, N., & Quarck, G. (2019). Impact of Galvanic Vestibular Stimulation on Anxiety Level in Young Adults. Frontiers in Systems Neuroscience, 13. 10.3389/fnsys.2019.00014

Pavani, F., Spence, C., & Driver, J. (2000). Visual Capture of Touch: Out-of-the-Body Experiences With Rubber Gloves. Psychological Science, 11(5), 353–359. 10.1111/1467-9280.00270

Pelagatti, C., Blini, E., & Vannucci, M. (2025). Catching Mind Wandering With Pupillometry: Conceptual and Methodological Challenges. WIREs Cognitive Science, 16(1), e1695. 10.1002/wcs.1695

Pervaz, I., Thurn, L., Vezzani, C., Kaluza, L., Kühnel, A., & Kroemer, N. B. (2025). Does transcutaneous auricular vagus nerve stimulation alter pupil dilation? A living Bayesian meta-analysis. Brain Stimulation, 18(2), 148–157. 10.1016/j.brs.2025.01.022

Pfeiffer, C., Serino, A., & Blanke, O. (2014). The vestibular system: A spatial reference for bodily self-consciousness. Frontiers in Integrative Neuroscience, 8, 31. 10.3389/fnint.2014.00031

Pilarczyk, J., Janeczko, W., Sterna, R., & Kuniecki, M. (2021). Are emotional objects visually salient? The Emotional Maps Database. Journal of Visual Communication and Image Representation, 79, 103221. 10.1016/j.jvcir.2021.103221

Pliego, A., Soto, E., Pliego, A., & Soto, E. (2025). Galvanic Vestibular Stimulation and Its Effects on Sympathetic Nervous System Activation. Journal of Integrative Neuroscience, 24(11). 10.31083/JIN45042

Pocock, S. J. (1977). Group sequential methods in the design and analysis of clinical trials. Biometrika, 64(2), 191–199. 10.1093/biomet/64.2.191

Ponzo, S., Kirsch, L. P., Fotopoulou, A., & Jenkinson, P. M. (2018). Balancing body ownership: Visual capture of proprioception and affectivity during vestibular stimulation. Neuropsychologia, 117, 311–321. 10.1016/j.neuropsychologia.2018.06.020

Preuss, N., Ellis, A. W., & Mast, F. W. (2015). Negative emotional stimuli enhance vestibular processing. Emotion, 15(4), 411–415. 10.1037/emo0000092

Preuss, N., Hasler, G., & Mast, F. W. (2014). Caloric Vestibular Stimulation Modulates Affective Control and Mood. Brain Stimulation, 7(1), 133–140. 10.1016/j.brs.2013.09.003

Quadt, L., Critchley, H. D., & Garfinkel, S. N. (2018). The neurobiology of interoception in health and disease. Annals of the New York Academy of Sciences, 1428(1), 112–128. 10.1111/nyas.13915

Quinn, V. F., MacDougall, H. G., & Colagiuri, B. (2015). Galvanic Vestibular Stimulation: A new model of placebo-induced nausea. Journal of Psychosomatic Research, 78(5), 484–488. 10.1016/j.jpsychores.2014.12.011

R Core Team. (2023). *R: A Language and Environment for Statistical Computing*. R Foundation for Statistical Computing. https://www.R-project.org/

Reynaud, A. J., Blini, E., Koun, E., Macaluso, E., Meunier, M., & Hadj-Bouziane, F. (2021). Atomoxetine modulates the contribution of low-level signals during free viewing of natural images in rhesus monkeys. Neuropharmacology, 182, 108377. 10.1016/j.neuropharm.2020.108377

Rode, G., Charles, N., Perenin, M. T., Vighetto, A., Trillet, M., & Aimard, G. (1992). Partial remission of hemiplegia and somatoparaphrenia through vestibular stimulation in a case of unilateral neglect. Cortex; a Journal Devoted to the Study of the Nervous System and Behavior, 28(2), 203–208.

Rode, G., Perenin, M. T., & Boisson, D. (1995). [Neglect of the representational space: Demonstration by mental evocation of the map of France]. Revue neurologique, 151(3), 161–164.

Seth, A. K. (2013). Interoceptive inference, emotion, and the embodied self. Trends in Cognitive Sciences, 17(11), 565–573. 10.1016/j.tics.2013.09.007

Sharon, O., Fahoum, F., & Nir, Y. (2021). Transcutaneous Vagus Nerve Stimulation in Humans Induces Pupil Dilation and Attenuates Alpha Oscillations. The Journal of Neuroscience, 41(2), 320–330. 10.1523/JNEUROSCI.1361-20.2020

Singmann, H., Bolker, B., Westfall, J., Aust, F., & Ben-Shachar, M. S. (2023). *afex: Analysis of Factorial Experiments*. https://CRAN.R-project.org/package=afex

Strauch, C., Wang, C.-A., Einhäuser, W., Van der Stigchel, S., & Naber, M. (2022). Pupillometry as an integrated readout of distinct attentional networks. Trends in Neurosciences, 45(8), 635–647. 10.1016/j.tins.2022.05.003

Toussaint, B., Heinzle, J., & Stephan, K. E. (2024). A computationally informed distinction of interoception and exteroception. Neuroscience & Biobehavioral Reviews, 159, 105608. 10.1016/j.neubiorev.2024.105608

Tsakiris, M. (2017). The multisensory basis of the self: From body to identity to others. Quarterly Journal of Experimental Psychology, 70(4), 597–609. 10.1080/17470218.2016.1181768

Tsakiris, M., & Critchley, H. (2016). Interoception beyond homeostasis: Affect, cognition and mental health. Philosophical Transactions of the Royal Society B: Biological Sciences, 371(1708), 20160002. 10.1098/rstb.2016.0002

Utz, K. S., Korluss, K., Schmidt, L., Rosenthal, A., Oppenländer, K., Keller, I., & Kerkhoff, G. (2011). Minor adverse effects of galvanic vestibular stimulation in persons with stroke and healthy individuals. Brain Injury, 25(11), 1058–1069. 10.3109/02699052.2011.607789

Voeten, C. C. (2019). Using ‘buildmer’to automatically find & compare maximal (mixed) models. R Package Version, 1(6), 1–7.

Wickham, H. (2016). ggplot2: Elegant Graphics for Data Analysis. Springer.

Wickham, H., François, R., Henry, L., Müller, K., & Vaughan, D. (2023). *dplyr: A Grammar of Data Manipulation*. https://CRAN.R-project.org/package=dplyr

Yates, B. J., Bolton, P. S., & Macefield, V. G. (2014). Vestibulo-Sympathetic Responses. Comprehensive Physiology, 4(2), 851–887. 10.1002/j.2040-4603.2014.tb00563.x

Zeller, D., Litvak, V., Friston, K. J., & Classen, J. (2015). Sensory Processing and the Rubber Hand Illusion—An Evoked Potentials Study. Journal of Cognitive Neuroscience, 27(3), 573–582. 10.1162/jocn_a_00705

zu Eulenburg, P., Caspers, S., Roski, C., & Eickhoff, S. B. (2012). Meta-analytical definition and functional connectivity of the human vestibular cortex. NeuroImage, 60(1), 162–169. 10.1016/j.neuroimage.2011.12.032

