## Supplementary 1 for "Nausea reshapes visual exploration and pupillary responses to affective images"

### Power Analysis and Sampling Plan

Supplementary File 1

#### Setup

We need the following libraries.

```
#libraries
library("pwr")
library("rpact")
library("gsDesign")
library("ggplot2")
```

#### Background

Ponzo et al. (2018, *Neuropsychologia*) analyzed 23 participants (after the exclusions). In their study, GVS (both left- and right- anodal conditions) and SHAM stimulations were administered in the context of the rubber hand illusion paradigm. The main outcome was the proprioceptive drift, which provides an objective proxy of visual capture during vestibular perturbances. The hemispheric effect of GVS (the difference between L- and R-GVS, L-GVS increasing proprioceptive drift and thus visual capture) was associated with a large effect size (d= 1.08, which roughly corresponds to a partial eta squared well above 0.2). Note that the contrast refers to the difference between L-GVS and R-GVS conditions, but SHAM was associated with a pattern very similar to R-GVS (not tested statistically). Also note that Ferré et al. (2015, *Neuropsychologia*) obtained similar results in terms of summary statistics (the partial eta squared was about 0.16)… Except that the pattern was reversed and L-GVS *decreased* visual capture. Our design, however, introduces another factor: valence, that is the category of images that are presented on screen (3 levels). Notwithstanding the several, important differences between the rubber hand paradigm and our setting, this factor may be somewhat akin to "speed" in Ponzo and colleagues. Speed (fast or slow) was meant to manipulate the affective nature of touch (CT-optimal or not). Hence, the three-way interaction between stimulation, synchrony, and speed is probably the closest effect to our experimental design. The effect size of this interaction was pes = 0.193.

For all subsequent analyses, this index is better transformed into Cohen's *f2*.

```
#pes to f2
f2= 0.193/(1 - 0.193)
f2
```

```
## [1] 0.2391574
```

What is the sample size required in order to have at least 90% power?

```
pwr.f2.test(u= 2,
            v= NULL, #find N, aka denominator dfs
            f2= f2,
            sig.level= 0.05,
            power= 0.9)$v
```

```
## [1] 53.028
```

```
53.028/2 + 1 #from dfs to N
```

```
## [1] 27.514
```

That would be 28 participants at minimum.

A few key specifications:

. We are looking for the three-way interaction between Image Category (Positive, Neutral, Negative) x Session (GVS, SHAM) x ( Stimulation (ON, OFF). Numerator degrees of freedom is therefore u= (2-1)(2-1)(3-1)= 2. An argument could be made about having Stimulation type as one only variable which would have 4 levels, thereby increasing slightly this parameter to u= 6. We choose Dfn= 2 for conservative reasons.

. We have estimated power for a general f test. The assumption is that different images within category will be collapsed. LMEMs with crossed random effects are much better suited (and powerful) for this case but require a set of additional assumptions for which we don't really have an informed starting point.

#### Sensitivity analysis

We estimate statistical power for a range of sample and effect sizes, striving for a more cautious sampling design.

```
#range of target sample sizes
SS= 30:42

#range of effect sizes to test
#in partial eta squared
ES= seq(0.1, 0.2, by= 0.02)

#note: this is f, not f2
sqrt(range(ES)/(1 - range(ES)))
```

```
## [1] 0.3333333 0.5000000
```

```
#expand
DF= expand.grid(EffectSize= ES, #range given as partial eta squared
                SampleSize= SS)

#transform pes to f2
DF$f2= DF$EffectSize/(1 - DF$EffectSize)
DF$EffectSize= as.factor(DF$EffectSize)


#compute
DF$Power= pwr.f2.test(u= 2, #Numerator dfs
                      v= 2*(DF$SampleSize - 1), #denominator dfs
                      f2= DF$f2,
                      sig.level= 0.05,
                      NULL)$power
```

Depicted:

```r
ggplot(DF, aes(x= SampleSize, y= Power,
               group= EffectSize, color= EffectSize)) +
  theme_bw() +
  labs(x= "Sample Size", y= "Power", color= "Effect Size") +
  geom_hline(yintercept= c(0.8, 0.9, 0.95),
             linewidth= 1,
             color= c("light gray", "gray", "dark gray"),
             linetype= "dashed") +
  geom_smooth(method= "loess",
              se= F, linewidth= 1.15) +
  geom_point(shape=19, size=1.6) +
  xlim(c(28, 44)) +
  ylim(c(0.5, 1)) +
  scale_color_manual(
  values = heat.colors(length(ES)),
  breaks = rev(as.character(ES)),
  name = expression(paste("Effect Size: " * eta^2[p]))) +
  theme(
    text = element_text(
      size = 11,
      face = "bold",
      color = "black"
    ),
    axis.text.x = element_text(
      size = 11,
      face = "bold",
      color = "black"
    ),
    axis.text.y = element_text(
      size = 11,
      face = "bold",
      color = "black"
    )) +
  annotate("label", x= 30, y= 0.55,
           label= "Interim \nAnalysis", fontface= 2, size= 2.5) +
  annotate("label", x= 42, y= 0.55,
           label= "Final \nAnalysis", fontface= 2, size= 2.5) +
  ggtitle("Power Analysis") +
  theme(legend.box.margin = margin(-10, -5, -10, -10))
```

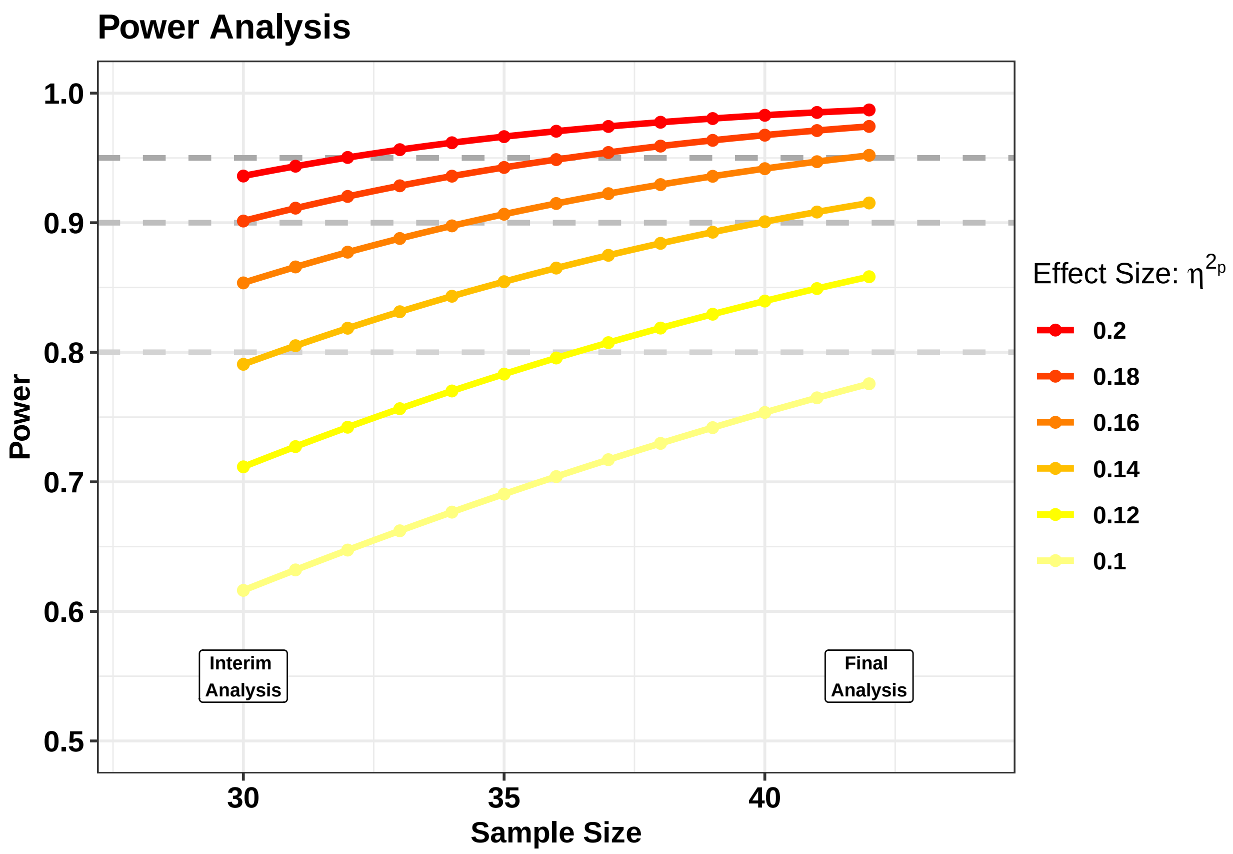


##### Sequential testing

In order to optimize data sampling and recruitment, we correct the alpha thresholds in order to preserve a cumulative 5% error rate across the experiment.

There are multiple ways (and libraries) to do it, we choose Pocock's correction for its simplicity. See: link.

```
seqDesign = gsDesign::gsDesign(
  k = 2,  #2 peeks
  test.type = 2,  #two-sided, symmetric
  alpha = 0.05,  #error rate
  sfu = "Pocock",  #discount function
  timing = c(30 / 42, 1)#unequally spaced
)

#boundaries
pnorm(-abs(seqDesign$upper$bound))
```

```
## [1] 0.03383435 0.03383435
```

Thus, we correct alpha to 0.034 and use it as our evidence threshold.

### Session info

```r
sessionInfo()
```

```
## R version 4.5.1 (2025-06-13 ucrt)
## Platform: x86_64-w64-mingw32/x64
## Running under: Windows 11 x64 (build 26200)
##
## Matrix products: default
##   LAPACK version 3.12.1
##
## locale:
## [1] LC_COLLATE=English_United States.utf8
## [2] LC_CTYPE=English_United States.utf8
## [3] LC_MONETARY=English_United States.utf8
## [4] LC_NUMERIC=C
## [5] LC_TIME=English_United States.utf8
##
## time zone: Europe/Rome
## tzcode source: internal
##
## attached base packages:
## [1] stats     graphics  grDevices utils     datasets  methods   base
##
## other attached packages:
## [1] ggplot2_4.0.1  gsDesign_3.8.0 rpact_4.3.0    pwr_1.3-0
##
## loaded via a namespace (and not attached):
##  [1] gt_1.3.0           rappdirs_0.3.4     generics_0.1.4     tidyr_1.3.2
##  [5] xml2_1.5.2         r2rtf_1.3.0        lattice_0.22-7     digest_0.6.39
##  [9] magrittr_2.0.4     evaluate_1.0.5     grid_4.5.1         RColorBrewer_1.1-3
## [13] fastmap_1.2.0      Matrix_1.7-3       mgcv_1.9-3         purrr_1.2.1
## [17] scales_1.4.0       textshaping_1.0.4  cli_3.6.5          rlang_1.1.7
## [21] splines_4.5.1      withr_3.0.2        yaml_2.3.12        otel_0.2.0
## [25] tools_4.5.1        dplyr_1.1.4        vctrs_0.7.0        R6_2.6.1
## [29] lifecycle_1.0.5    fs_1.6.6           ragg_1.5.0         pkgconfig_2.0.3
## [33] pillar_1.11.1      gtable_0.3.6       glue_1.8.0         Rcpp_1.1.1
## [37] systemfonts_1.3.1  xfun_0.56          tibble_3.3.1       tidyselect_1.2.1
## [41] rstudioapi_0.18.0  knitr_1.51         farver_2.1.2       xtable_1.8-4
## [45] htmltools_0.5.9    nlme_3.1-168       rmarkdown_2.30     labeling_0.4.3
## [49] compiler_4.5.1     S7_0.2.1
```
