## Supplementary 2 for "Nausea reshapes visual exploration and pupillary responses to affective images"

### Description of the final stimuli

Supplementary File 2

Table 1: Valence and Arousal Ratings per Stimulus

| Category | Theme | Valence | Arousal |
|---|---|---|---|
| Negative | Miserable pose 3 | 1.107843 | 5.148515 |
| Negative | Dead bodies 1 | 1.198020 | 4.514563 |
| Negative | Dead bodies 2 | 1.247525 | 4.475728 |
| Negative | Dead bodies 3 | 1.323529 | 5.128713 |
| Negative | KKK rally 1 | 1.343137 | 4.702970 |
| Negative | Tumor 1 | 1.401961 | 4.940594 |
| Negative | Scary face 1 | 1.777778 | 4.519231 |
| Negative | Frustrated pose 3 | 2.351852 | 3.557692 |
| Negative | Frustrated pose 1 | 2.352941 | 4.247525 |
| Negative | Angry face 3 | 2.500000 | 3.663366 |
| Negative | Intensity 1 | 2.813725 | 4.148515 |
| Negative | Angry pose 1 | 2.814815 | 3.615385 |
| Neutral | Surgery 3 | 3.675926 | 3.913461 |
| Neutral | Boxing 2 | 3.696078 | 4.198020 |
| Neutral | Neutral face 4 | 3.891089 | 2.178218 |
| Neutral | Couple 3 | 4.009259 | 3.442308 |
| Neutral | Neutral face 3 | 4.504950 | 2.621359 |
| Neutral | School 5 | 4.539216 | 2.633663 |
| Neutral | Picnic 3 | 4.546296 | 3.269231 |
| Neutral | Doctor 1 | 4.666667 | 3.148515 |
| Neutral | Neutral face 2 | 4.675926 | 2.182692 |
| Neutral | Neutral face 5 | 4.980392 | 3.772277 |
| Neutral | Neutral pose 1 | 5.058823 | 3.019802 |
| Neutral | Swimming 1 | 5.059406 | 3.621359 |
| Positive | Nude couple 9 | 5.166667 | 5.059406 |
| Positive | Skydiving 2 | 5.333333 | 4.603960 |
| Positive | Nude couple 2 | 5.441177 | 4.405941 |
| Positive | Rafting 4 | 5.583333 | 4.048077 |
| Positive | Couple 6 | 5.607843 | 4.138614 |
| Positive | Smiling face 1 | 5.638889 | 4.125000 |
| Positive | Nude couple 5 | 5.685185 | 5.346154 |
| Positive | Yoga 4 | 5.725490 | 3.990099 |
| Positive | Couple 4 | 5.970588 | 4.534654 |
| Positive | Wedding 6 | 6.000000 | 3.922330 |
| Positive | Mother 7 | 6.039604 | 4.300971 |
| Positive | Mother 6 | 6.078431 | 4.445545 |

Table 2: Summary Statistics by Category (Mean ± SD)

| Category | Mean Valence | SD | Mean Arousal | SD |
|---|---|---|---|---|
| Negative | 1.85 | 0.67 | 4.39 | 0.56 |
| Neutral | 4.44 | 0.51 | 3.17 | 0.66 |
| Positive | 5.69 | 0.29 | 4.41 | 0.43 |

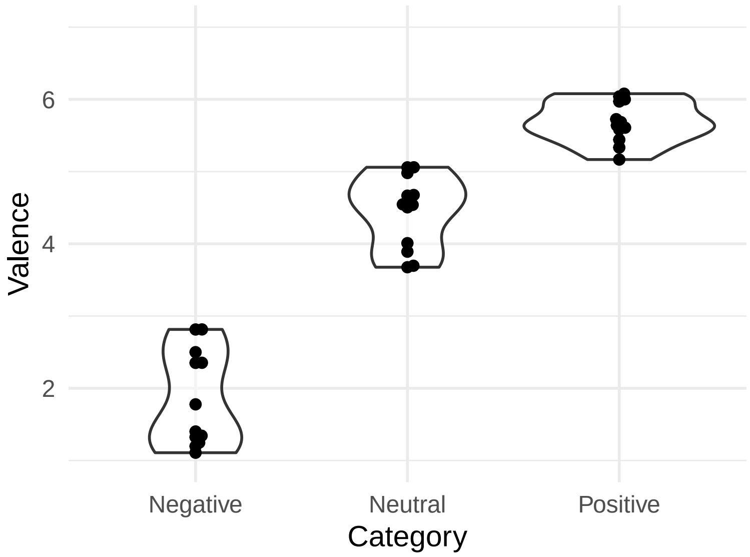


Figure 1: Valence Ratings

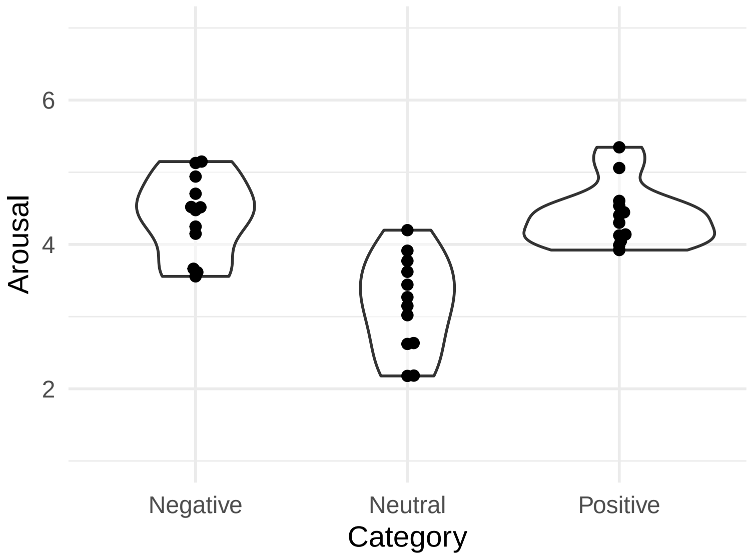


Figure 2: Arousal Ratings
