## Supplementary Figures for "Nausea reshapes visual exploration and pupillary responses to affective images"

#### Supplementary Figures 1 to 11

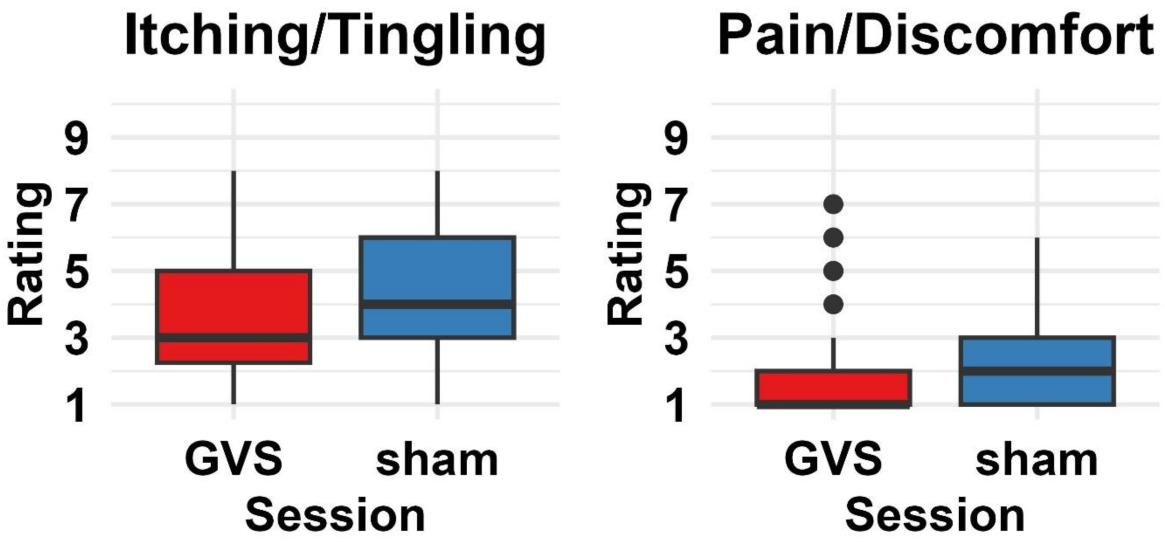


**Supplementary Figure 1. GVS and sham ratings – non vestibular sensations.** Non-specific itching and tingling sensations induced by the electrical stimulation did not differ between conditions ($t_{(41)}$= -1.88, $p$= 0.067); if anything, itching was more intense during the sham session, showing that the control condition worked as intended in inducing arousal originating from the physical sensations elicited by the electrodes. When participants were asked to rate the overall pain and discomfort caused by the procedure, however, the two conditions were closely matched ($t_{(41)}$= -0.27, $p$= 0.79).

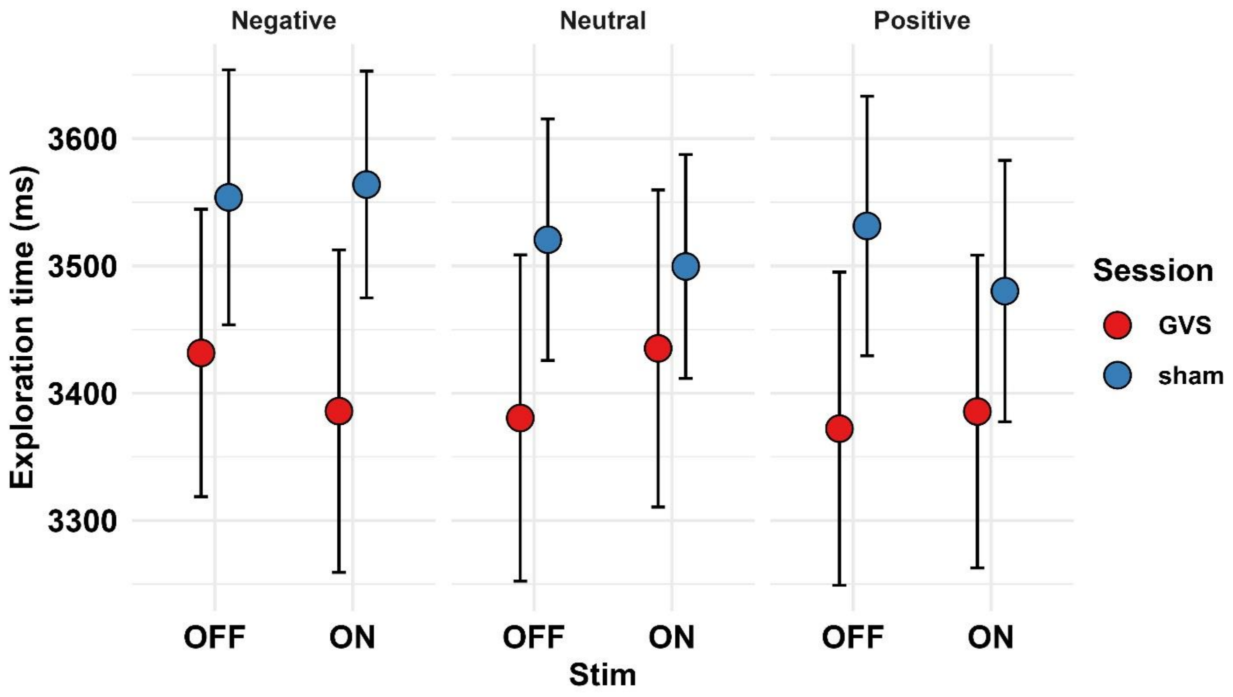


**Supplementary Figure 2. IVT: basic exploration parameters.** Total dwell time (mean ± SEM) did not differ across Sessions, Category, Stimulation, or Susceptibility to GVS. There was a visual trend for GVS ro reduce total dwell time, but the corresponding effect was not statistically significant (p= 0.224). There was large variability in this measure. We can decompose further dwell time into 1) number of fixations and 2) mean fixation duration. These variables are shown in the next supplementary figures.

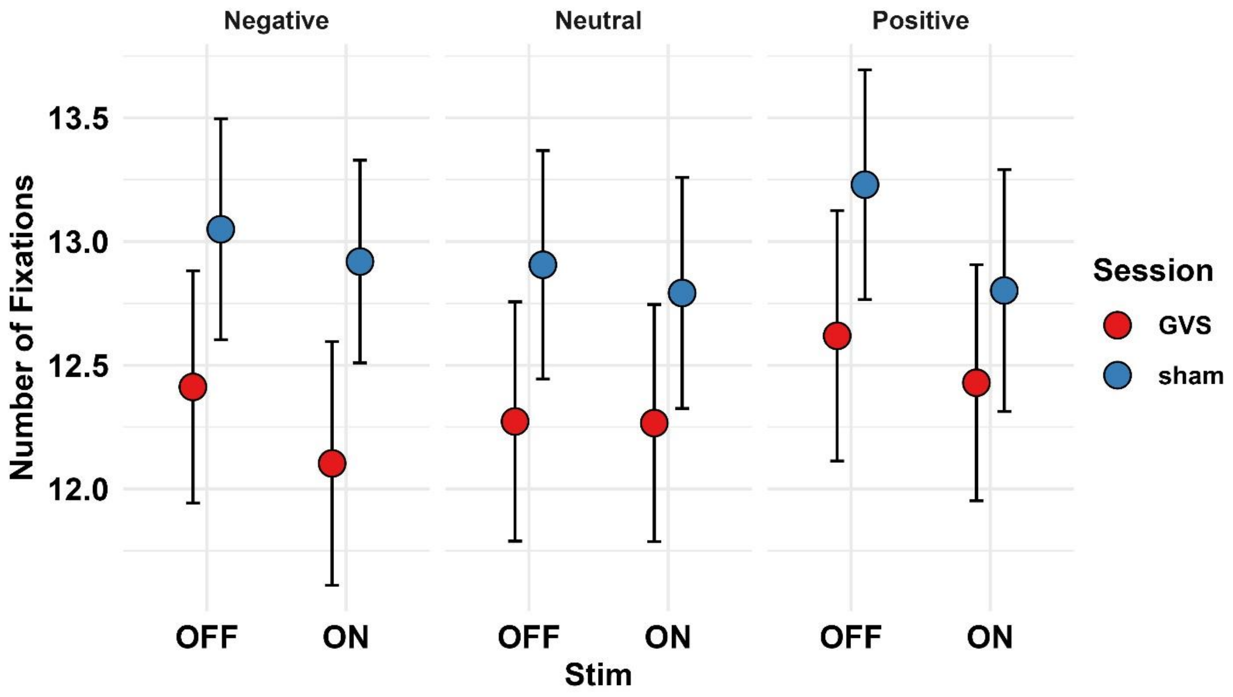


**Supplementary Figure 3. IVT: basic exploration parameters.** The average number of fixations (here shown mean ± SEM) was smaller for GVS (although not significantly so).

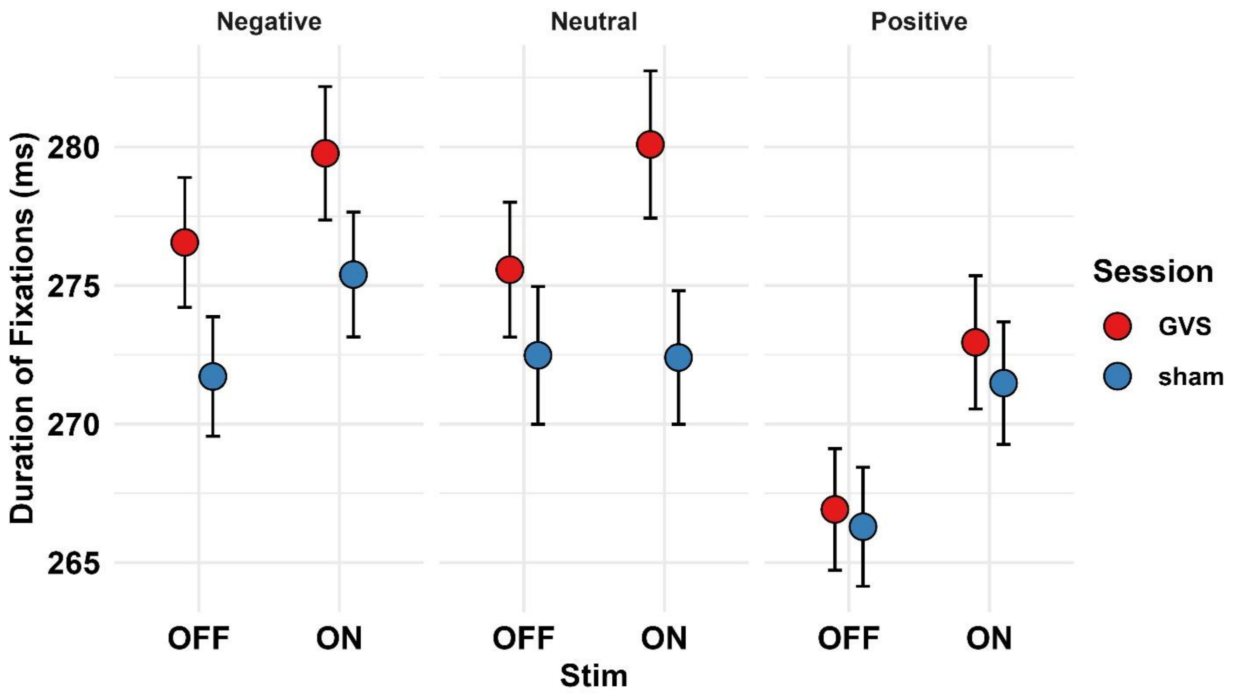


**Supplementary Figure 4. IVT: basic exploration parameters.** Each fixation (here mean ± SEM) was on average slightly longer during GVS. Again, differences were not statistically significant for the main effect of Session (GVS vs sham). However, there was a small tendency for electric stimulation (so both GVS and sham when online) to increase the average duration of fixation events (F (1, 35.94)= 3.29, p= 0.078). There was also a main effect of Category, with positive images being associated to shorter fixations overall.

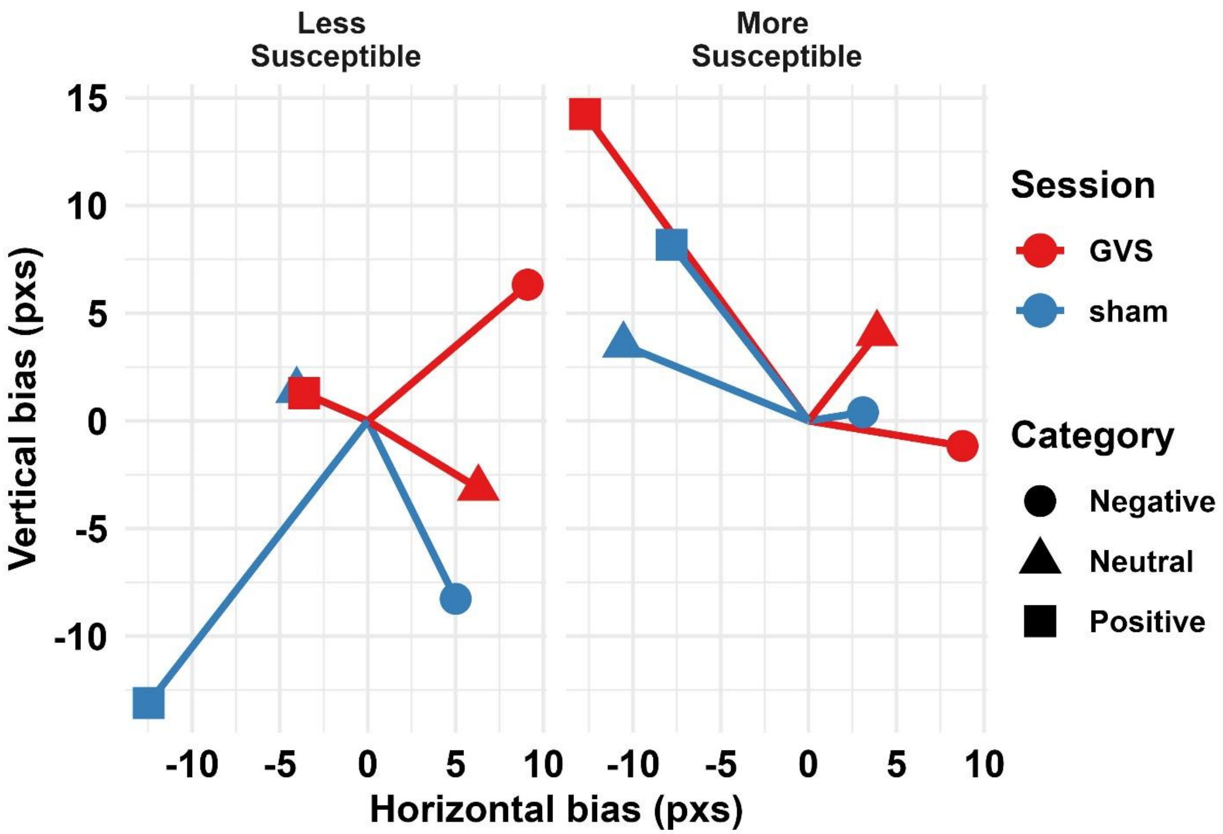


**Supplementary Figure 5. IVT: CoG displacements (ON-OFF).** This is an alternative depiction of Figure 3A in which the CoG *shifts* are depicted, for both the horizontal and vertical axes, in terms of differences between the respective online and offline stimulation condition. The plot shows the overall effects of GVS, which induces, on average, small rightward and upward biases, and the heterogeneity between different categories, thus showing that spatial biases are not fixed but depends to some extent on image content and valence.

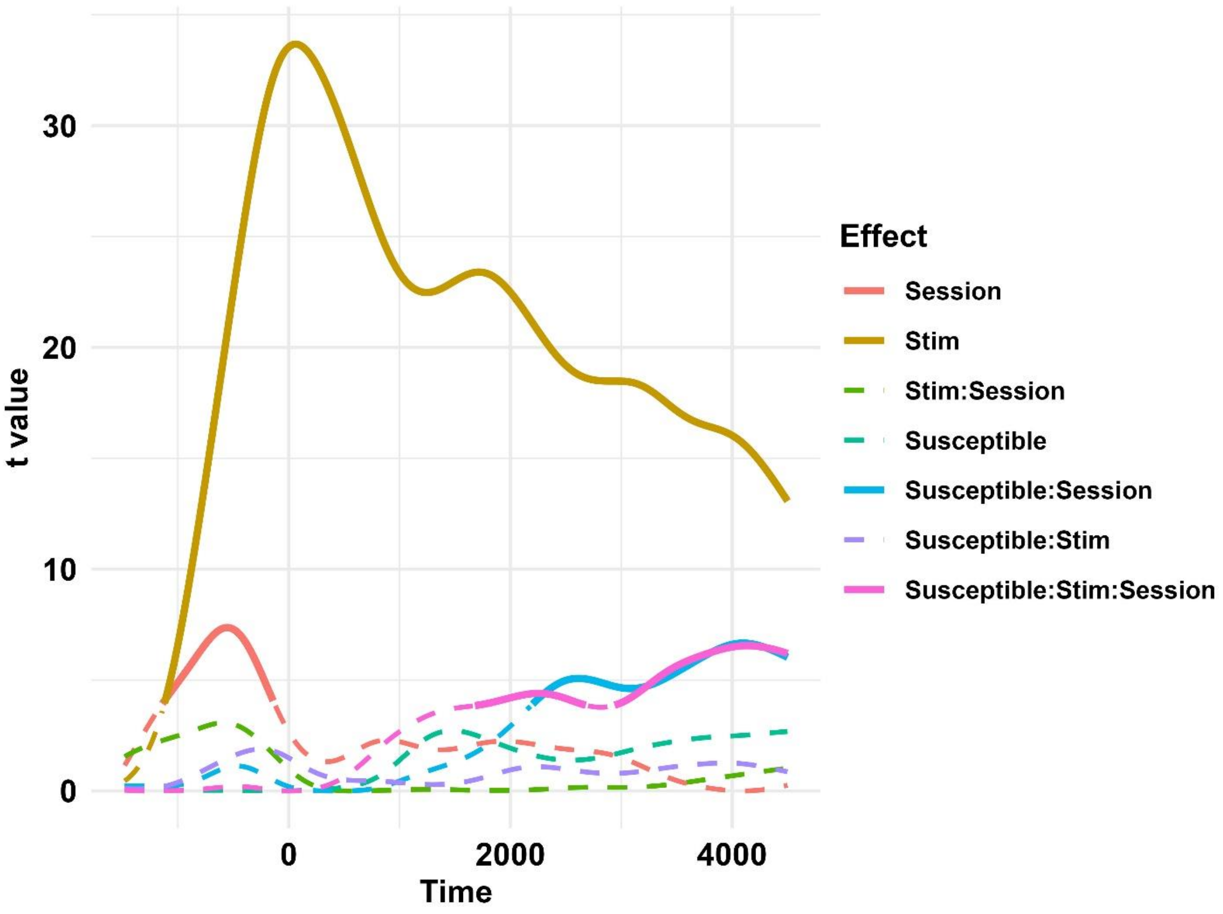


**Supplementary Figure 6. PT: LMEM modelling results**. This image depicts the statistics associated with LMEMs modelling. Linear Mixed Effects Models were computed for each timepoint; fixed factors were tested with Likelihood Ratio Tests. The plot shows the t statistics (y axis) for each timepoint (x axis). This analysis used a baseline prior to the electrical stimulation and was intended to assess the effects of Session, Stimulation, and Susceptibility. Lines represent for each factor, if solid, timepoints significant at the 0.05 threshold (here uncorrected). The electric stimulation had a strong effect on pupil size changes, but there were session effects as well (GVS being associated with larger increase) especially in susceptible participants.

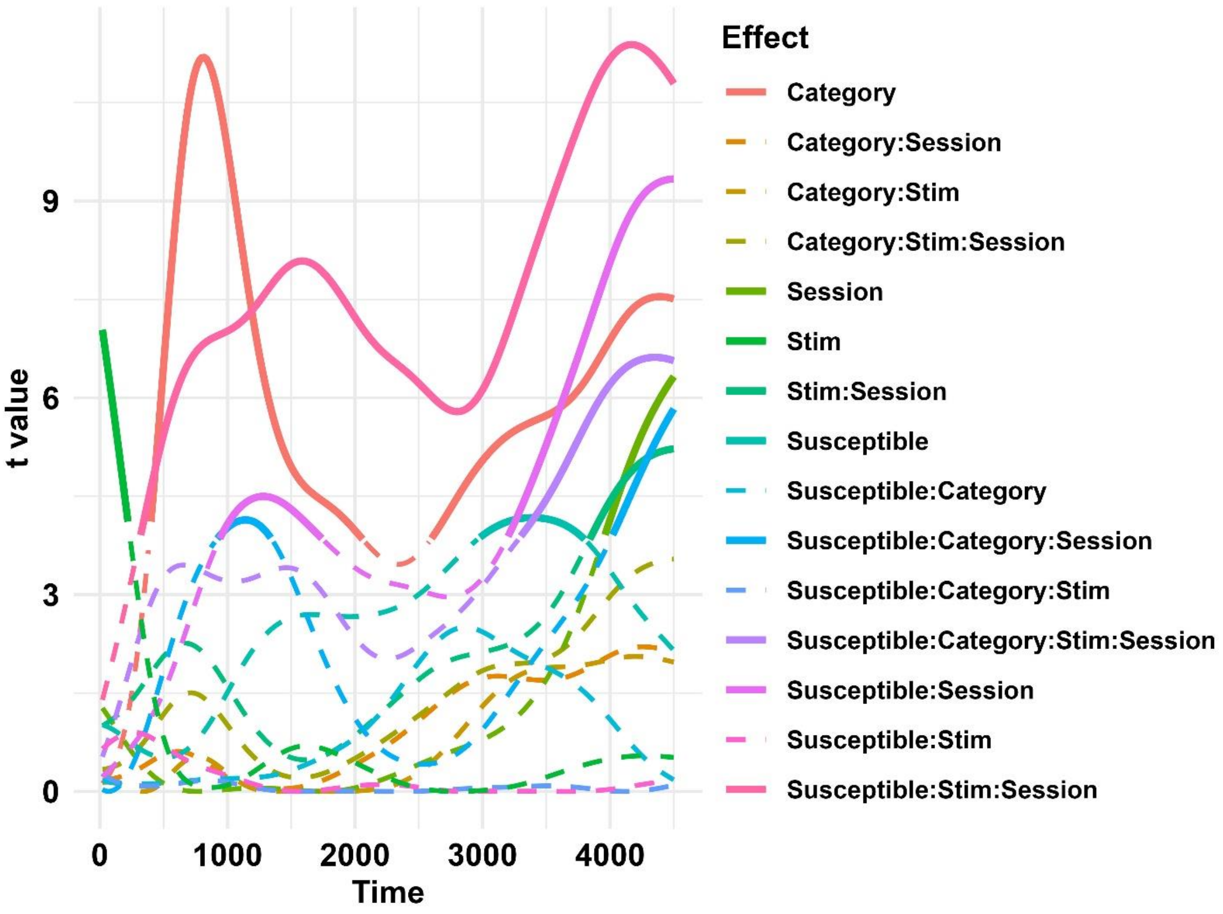


**Supplementary Figure 7. PT: LMEM modelling results**. This image depicts the statistics associated with LMEMs modelling. Linear Mixed Effects Models were computed for each timepoint; fixed factors were tested with Likelihood Ratio Tests. The plot shows the t statistics (y axis) for each timepoint (x axis). This analysis used a baseline prior to the presentation of images, and was intended to assess the effects of Category, Session, Stimulation, and Susceptibility. Lines represent for each factor, if solid, timepoints significant at the 0.05 threshold (here uncorrected). Image category had a strong effect on the pupils, with negatively-valenced images being associated with larger changes in pupil size. Multiple effects had a role in shaping the overall data pattern, most notably the 4-way interaction, discussed in the main text, suggesting that online GVS abolished, in highly susceptible participants, the effect of category outlined above.

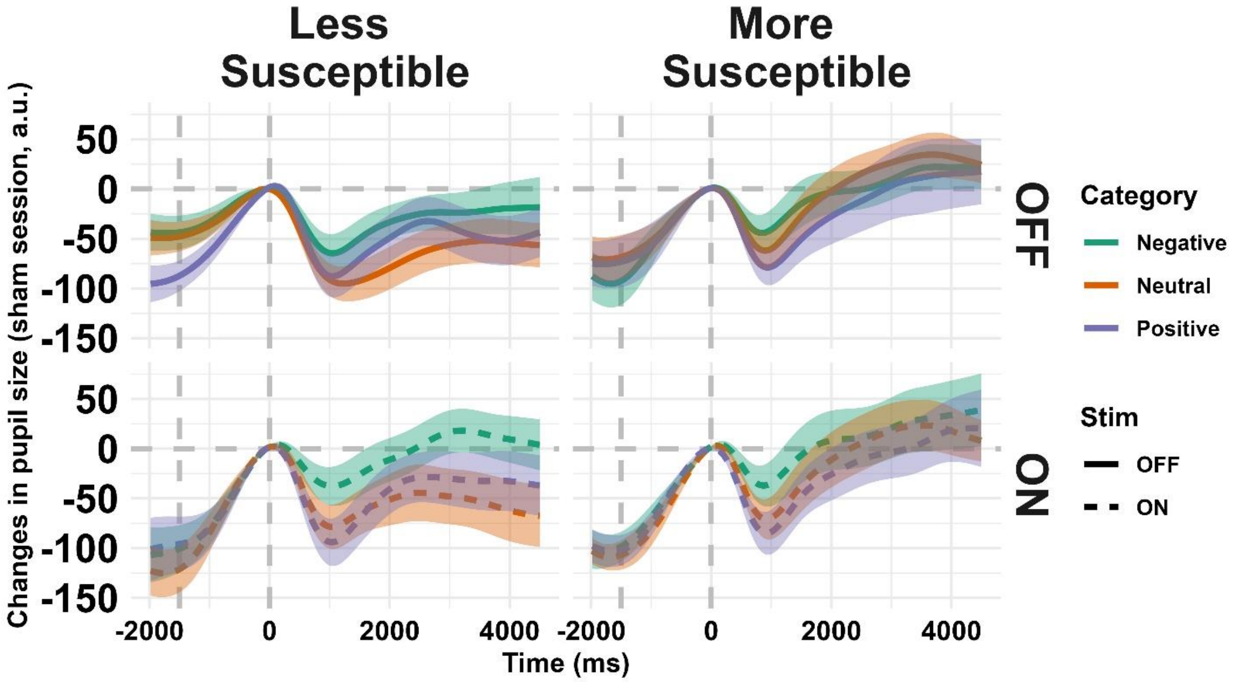


**Supplementary Figure 8. PT: phasic changes in pupil size, sham session.** Same as figure 5C but for the sham session.

### PT: phasic changes in pupil size, residuals

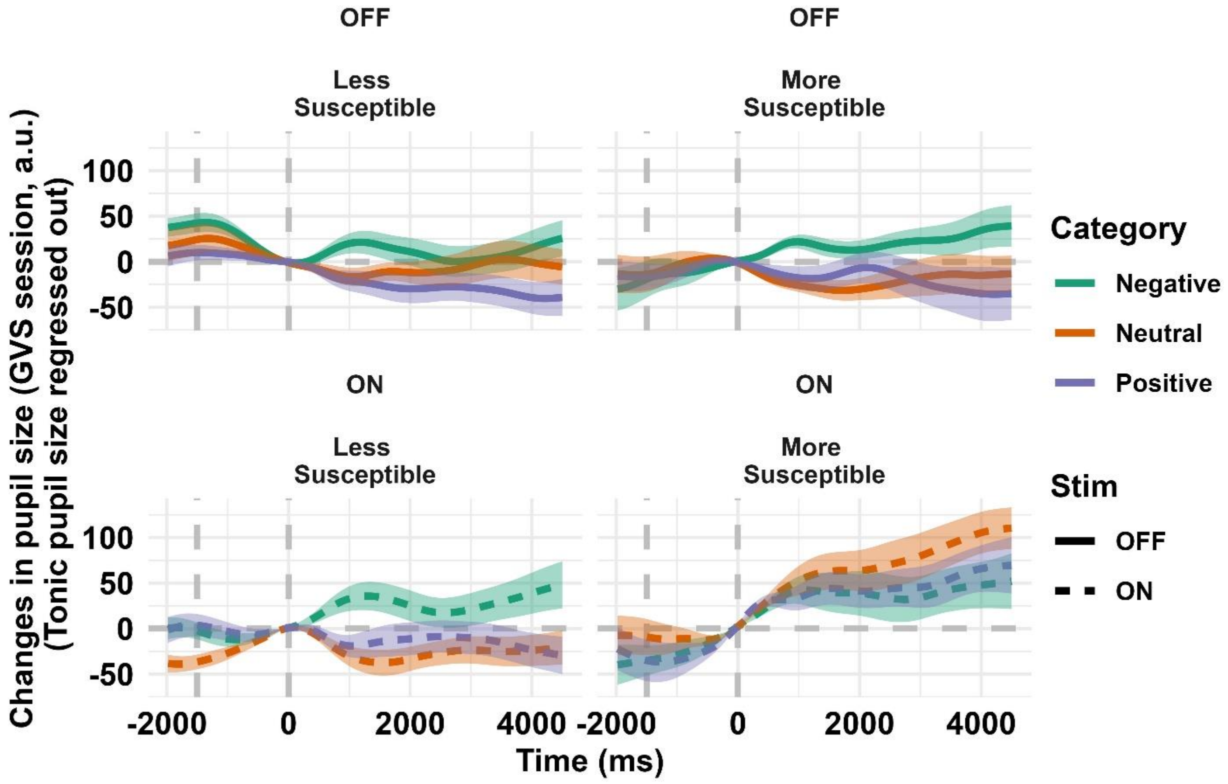


**Supplementary Figure 9. PT: phasic changes in pupil size, residuals.** Same as figure 5C, but here we regressed out from the data the (trial-wise) tonic pupil size. The size of the pupil was larger for susceptible participants during online GVS; however, the residuals consistently show a disrupted pattern of changes in pupil size for emotional images even when accounting for this factor.

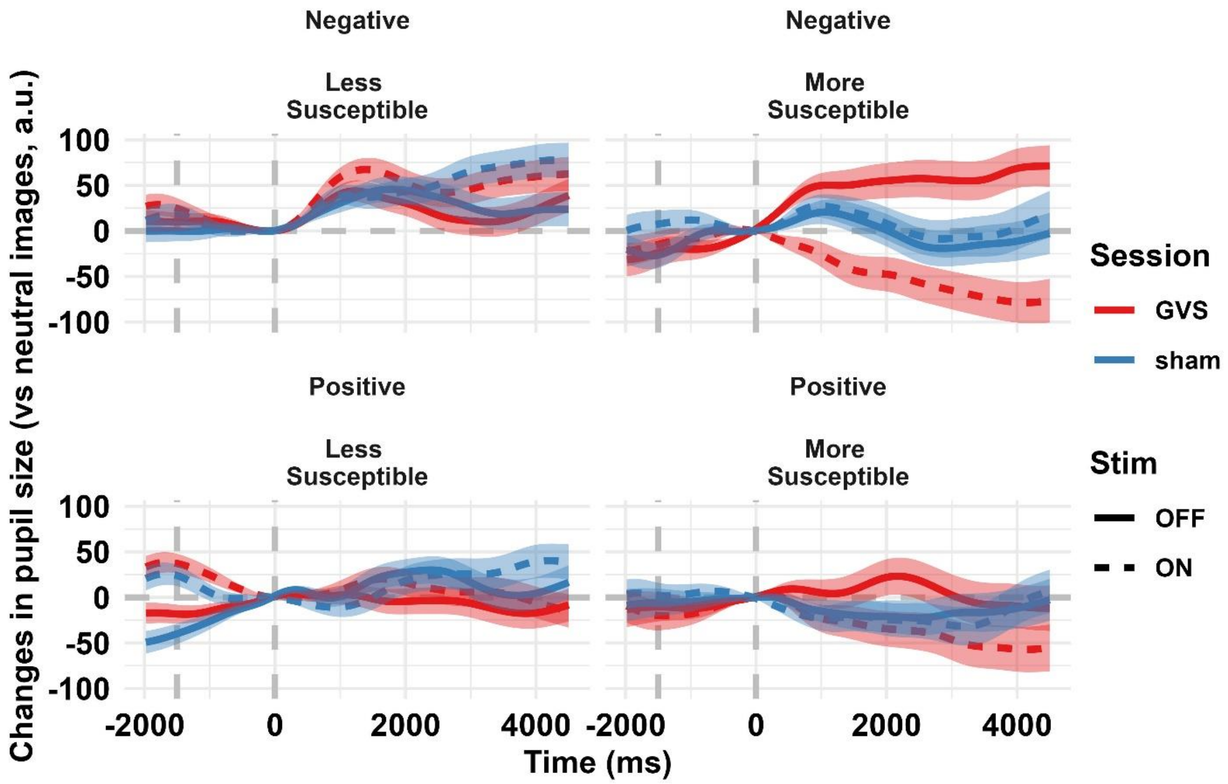


**Supplementary Figure 10. PT: phasic changes in pupil size, Δ changes from neutral.** Mean (SEM) changes in pupil size with respect to neutral images. Negative images were associated with pupil dilation. In less susceptible participants, this is shown as predominantly positive values in the image presentation phase. Susceptible participants, on the other hand, selectively for online GVS, presented an overall negative trend, meaning that negative images were rather associated with less dilation with respect to neutral images.

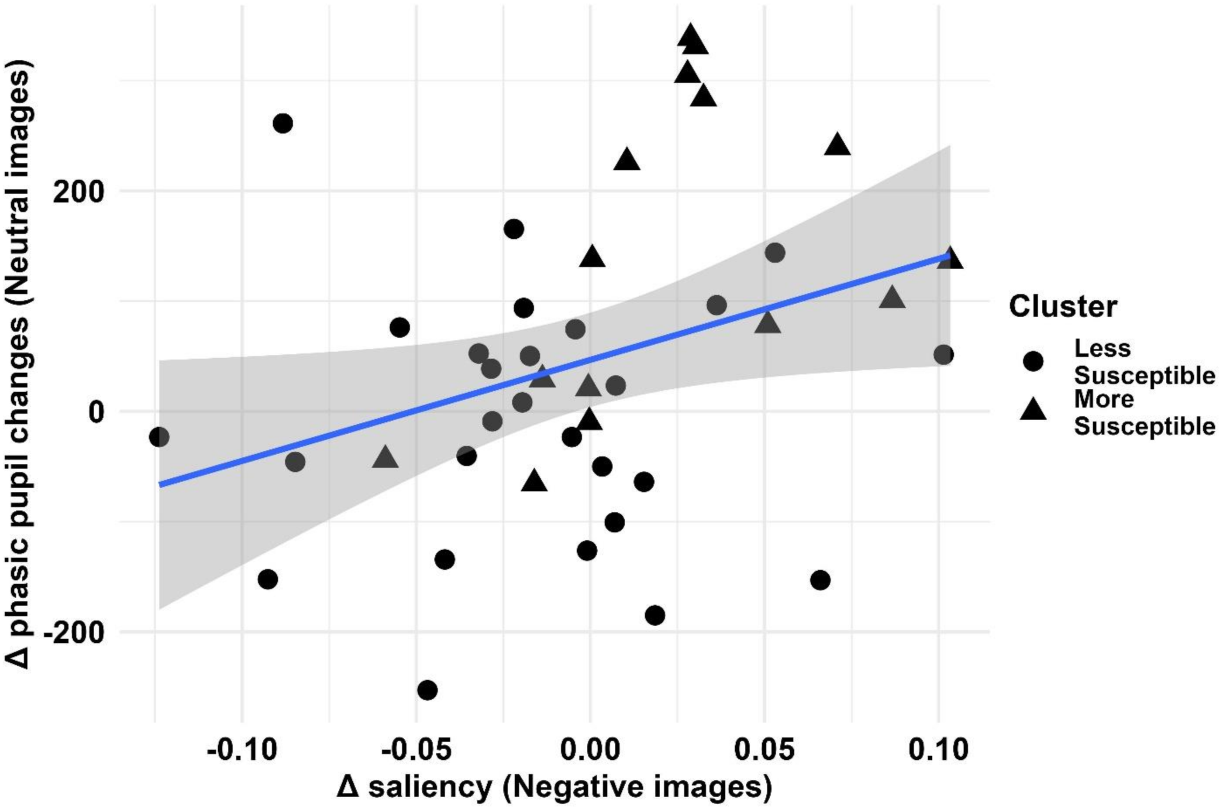


**Supplementary Figure 11. Correlation between pupillary dynamics and Δ saliency**. Altered phasic pupillary responses correlate with altered reliance on low-level features. We attempted to correlate the altered phasic responses with Δ saliency – that is the relative increase/decrease in reliance to low-level features from offline to online GVS observed for negative images in an independent task. This analysis is explorative: the correlation reported is seemingly very fragile and should therefore be interpreted with caution. We reported that changes in pupil size due to online GVS were much more pronounced, in highly susceptible participants, for neutral images. This GVS-specific gain correlates with Δ saliency for negative images ($r= 0.323$, $t(40)= 2.16$, $p= 0.037$). Participants who showed the largest pupil gain for neutral images also showed the largest increase in reliance on low-level visual features for negative images. It is likely that the positive correlation stems from susceptibility to GVS being a common defining factor. Still, the positive correlation strengthens the idea that Δ saliency may have been affected primarily due to a recalibration of affective responses to a new baseline. Results altogether suggest that vestibular input shifts the balance between emotion-driven salience and top-down inhibition, possibly via a modulation of lateralized autonomic pathways.
